# Revealing the polarity of actin filaments by cryo-electron tomography

**DOI:** 10.1101/2020.03.11.987263

**Authors:** Bruno Martins, Simona Sorrentino, Wen-Lu Chung, Meltem Tatli, Ohad Medalia, Matthias Eibauer

## Abstract

The actin cytoskeleton plays a fundamental role in numerous cellular processes, such as cell motility, cytokinesis, and adhesion to the extracellular matrix. Revealing the polarity of individual actin filaments in cells, would foster an unprecedented understanding of cytoskeletal processes and their associated mechanical forces. Cryo-electron tomography provides the means for high-resolution structural imaging of cells. However, the low signal-to-noise ratio of cryo-tomograms obscures the high frequencies and therefore the polarity of actin filaments cannot be directly measured. Here, we developed an approach that enables to determine the polarity of actin filaments in cellular cryo-tomograms. We applied it to reveal the actin polarity distribution in focal adhesions, and show a linear relation between actin polarity and distance from the apical boundary of the adhesion site.

## Introduction

Actin polymerization drives cell motility and is a central factor in mediating contractile forces in cells (Holmes et al., 1990; Merino et al., 2018; Pollard and Borisy, 2003). Actin filaments (Egelman et al., 1982; Galkin et al., 2015) assemble into complex networks (Malik-Garbi et al., 2019; Xu et al., 2012), which are essential for their activity in the cell. Reconstructing individual actin filaments at sub-nanometer resolution inside cells, would provide an unparalleled view on actin networks, and would allow a more fine-grained modelling of cytoskeletal based mechanical processes (Hervas-Raluy et al., 2019).

A prominent mechanosensitive process, involving actomyosin contractility, occurs at the integrin-based interaction sites of cells with the extracellular matrix (Burridge and Guilluy, 2016; Geiger et al., 2009). These interactions are mediated by adhesive structures such as focal adhesions (FAs) (Legate et al., 2011; Shemesh et al., 2005; Zaidel-Bar et al., 2007). The organization of proteins and the role of the actin network in FAs has been intensively studied using fluorescent and electron microscopy (Kanchanawong et al., 2010; Patla et al., 2010), however, the 3D architecture of FAs, including the polarity and position of each individual actin filament was not yet shown.

Cryo-electron tomography (cryo-ET) allows to reconstruct 3D density maps of unperturbed cells at a resolution of 2-3 nm (Beck and Baumeister, 2016; Lucic et al., 2005; Weber et al., 2019). Therefore, single actin filaments can readily be detected in tomograms of eukaryotic cells (Jasnin et al., 2019; Medalia et al., 2002). Moreover, using direct electron detectors permit to recognize the characteristic helical shape of F-actin that is in clear distinction to other cellular filaments. However, cryo-tomograms suffer from a low signal-to-noise ratio (SNR) (Forster et al., 2008; Pei et al., 2016), and from missing information due to the limited tilt range during data acquisition, referred to as the missing wedge (Lucic et al., 2005). Subtomogram averaging can compensate for these issues (Beck and Baumeister, 2016; Forster et al., 2005; Himes and Zhang, 2018; Schur et al., 2016), but it is computationally more demanding and is lacking the robustness and level of standardization as the procedures used for data analysis in single particle cryo-electron microscopy (Nogales, 2016; Scheres, 2012).

Here we developed a set of MATLAB scripts that enable, in conjunction with RELION (Scheres, 2012), the 3D reconstruction of actin filaments from cryo-tomograms. It is based on transforming subtomogram averaging into a single particle task, and the determination of their polarity is built on a robust statistical analysis of the individual filaments. Furthermore, this actin polarity toolbox (APT) features tools for spatial and topological analysis of the reconstructed actin filament networks and their visualization.

Using a correlative fluorescence microscopy and cryo-ET approach (Elad et al., 2013; Patla et al., 2010; Sartori et al., 2007), we unveil the 3D architecture of the actin cytoskeleton at FAs. We show that the actin polarity distribution correlates with the position along the FA and that regions of mixed polarity are concentrated at the periphery of the characteristic actin bundles.

## Results

### Determining actin polarity by APT

The first aim of APT is the reconstruction of an actin filament at a resolution that allows the unambiguous determination of the filament polarity (better than 20 Å). Therefore, APT requires cryo-tomograms and segmentations of the actin cytoskeleton as input data (Fig. 1a). Subsequently, the segmented filaments are subdivided into segments with equidistant spacing. We make use of the observation that a projection of a subtomogram along the electron beam axis is approximately invariant of the missing wedge orientation and can be treated as a single particle image (Supplemental Fig. 1). Thus, single particle image processing packages, such as cryoSPARC (Punjani et al., 2017) or RELION (Scheres, 2012), can be used to reconstruct the actin filament from the segments. Interestingly, this approach is not only restricted to filamentous structures (Supplemental Fig. 2).

**Figure 1.**
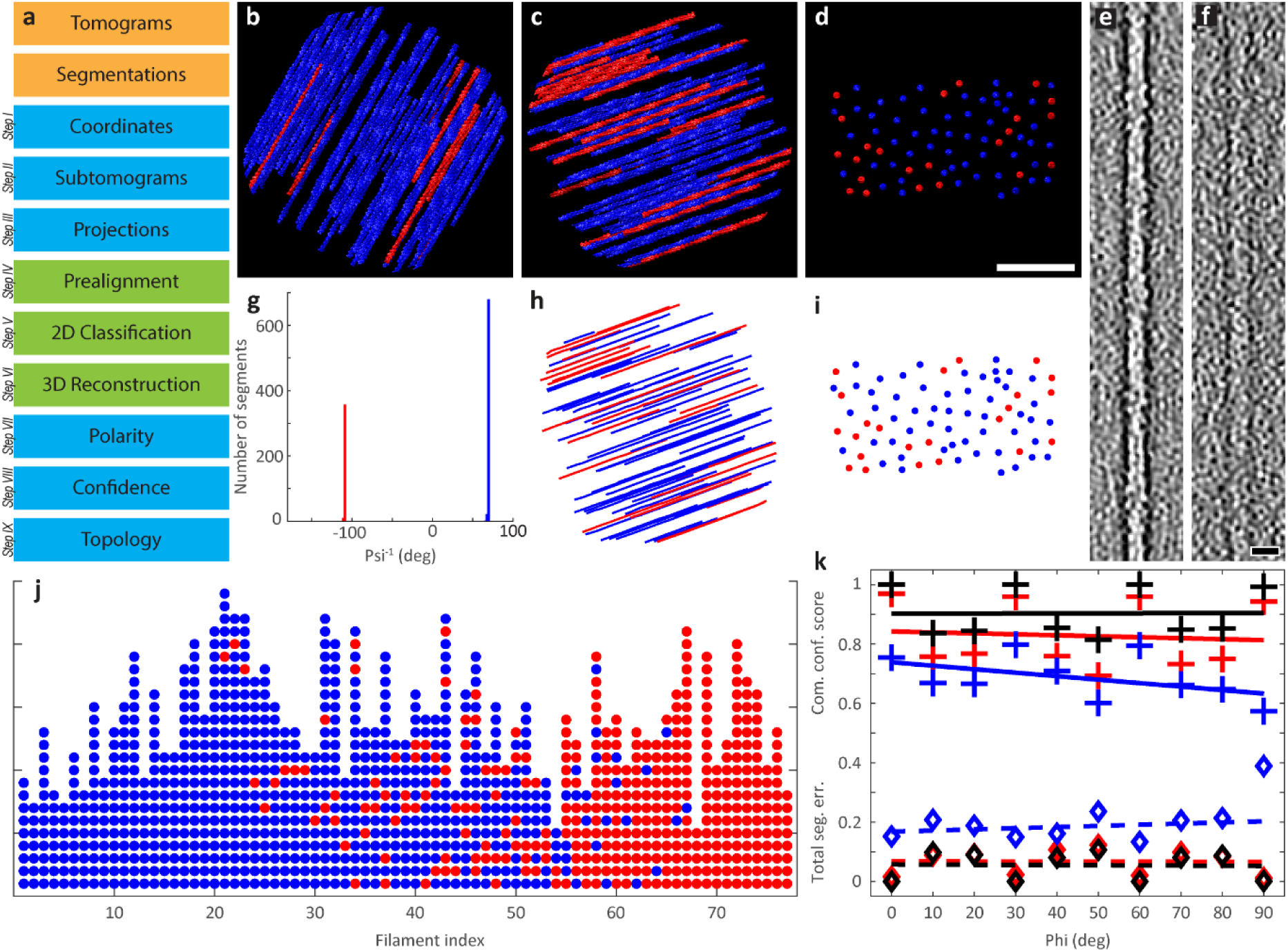
Polarity determination of modelled actin bundles. (**a**) The workflow of APT consists of consecutive modules. The orange modules symbolize the required input data. The blue modules were implemented in MATLAB, and the green modules were executed in RELION. (**b**) Top view of one of the modelled actin bundles, which were used for the validation of APT. The bundles were rendered without noise for visualization purposes. Here the angle *φ* between bundle and tilt-axis is 30°, and the fraction *ρ*_*r*_ of actin filaments, that are oriented in opposite direction (red colored filaments) was set to 0.1. (**c**) Top view of an additional bundle with *φ* increased to 70°, and *ρ*_*r*_ to 0.3. (**d**) The same bundle seen from the side with the viewing axis adjusted parallel to the filaments. The distance between the filaments is 10-37 nm. Scale bar 100 nm. (**e**) In order to quantify the impact of noise on the precision of APT, the bundles were modelled with defined SNRs. The depicted bandpass filtered slice of a filament was extracted from a bundle with SNR = 0.001. For comparison, the filament shown in (**f**) originates from a bundle with SNR = 0.0001. Scale bar 10 nm. (**g**) The modelled bundles were processed with APT. The plot shows the resulting *ψ*^−1^ histogram of the bundle displayed in (**c**). Segments in the blue peak mainly originate from blue filaments, and the opposite for the red peak. The filaments point with their plus-ends in opposite directions, therefore the peaks are separated by 180°. Since *ρ*_*r*_ was set to 0.3 in this bundle, the number of segments in the red peak is ≈30% of the total number of segments. (**h**) The line plot shows the filament positions and polarities of the bundle, shown in (**c**), as recovered by APT. It was affected with a SNR = 0.001 in this case, however, the match between original and recovered bundle (the side view is shown in (**i**), and can be compared to (**d**)) is almost error free. (**j**) Here the 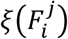 vectors of the recovered bundle are depicted. Segments, that are originating from the same filament are plotted as columns of circles. The colour scheme reflects to which peak the segments were assigned in the *ψ*^−1^ histogram, shown in (**g**), and the 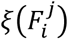 vectors are sorted according to their fraction of segments linked to the blue peak. In this representation of the filaments, segments with incorrect orientation determination appear as distortions of otherwise uniformly colored columns. (**k**) Three datasets of ten modelled actin bundles each, with SNRs of 0.01, 0.001, and 0.0001, were produced, subsequently processed with APT, and finally the ccs values and errors in the polarity determination were analyzed. Each cross marks the average of all ccs values in a bundle. The SNR can be identified by the color scheme: black, SNR = 0.01, red, SNR = 0.001, and blue, SNR=0.0001, respectively. A regression line was fitted for each SNR condition, and show that there is no trend to lower ccs values with increasing *φ* for SNRs ≳ 0.001 (black and red lines). Thus, the missing wedge induced anisotropy, which is proportional to *φ*, can be neglected for those SNR levels. However, for SNRs ≲ 0.0001 a correlation can be detected (blue line). Next, we calculated the average total segment error for each bundle, plotted as diamond symbols. Respective regression lines are delineated as dashed lines. Even for the markedly challenging SNR = 0.0001 the total segment error rarely exceeds 20%. Because APT uses a majority criterion for final polarity determination of a whole filament, segment errors <50% can be compensated.

**Figure 2.**
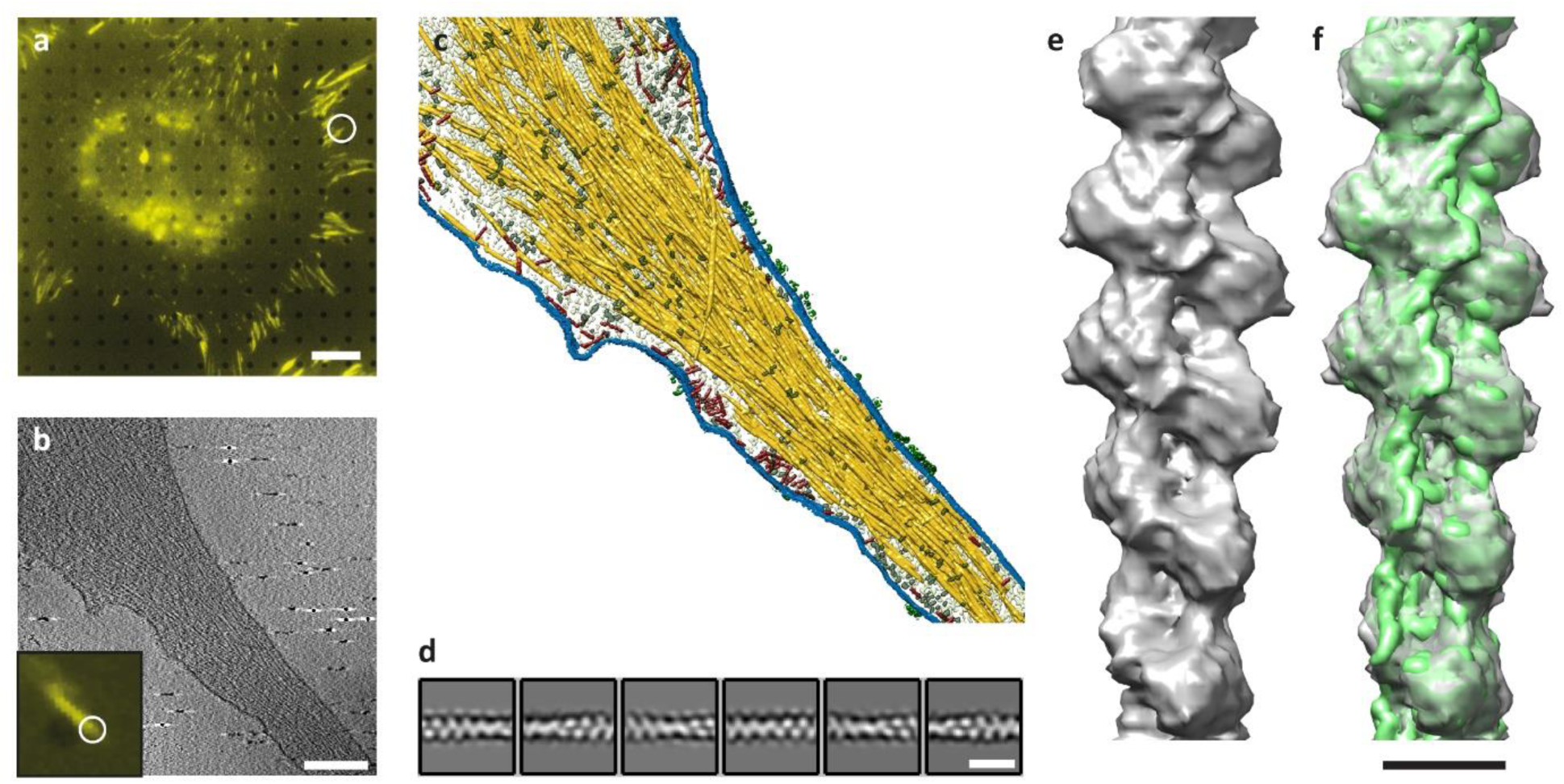
Cryo-tomography of FAs and actin filament structure from inside cells. Correlated microscopy combining fluorescence microscopy (**a, b**, inset) and cryo-ET (**b**) was used to identify FA sites. Scale bar in (**a**) 10 µm. (**b**) A 1.4 nm thick slice through a cryo-tomogram of a FA (**a, b**, white circles), shows individual actin filaments and plasma membrane. Scale bar 200 nm. (**c**) Surface rendering view of the FA site. Actin is depicted in yellow, membranes in blue, macromolecules in red and grey, and receptor densities in green. The segmentation of the actin filaments served as input for APT. (**d**) Class averages obtained by 2D classification of the extended dataset. Scale bar 18 nm. (**e**) Structure of an actin filament at FAs (Supplementary Fig. 9), shown as grey isosurface. (**f**) The in-vitro structure EMD-6179 (Galkin et al., 2015) (green isosurface) was docked into the in-situ structure. Scale bar 5 nm.

Our main goal is to determine the polarity of all the initially segmented filaments. Therefore, based on the filament average, APT calculates for each segment the position of its plus-end in the cryo-tomograms (Supplemental Fig. 3a). During the 3D reconstruction step, APT obscures the correlation between segments and filaments, thus for each filament multiple independent polarity observations can be statistically evaluated, which greatly reduces the error of polarity determination, as shown in the validation of the method (see below).

**Figure 3.**
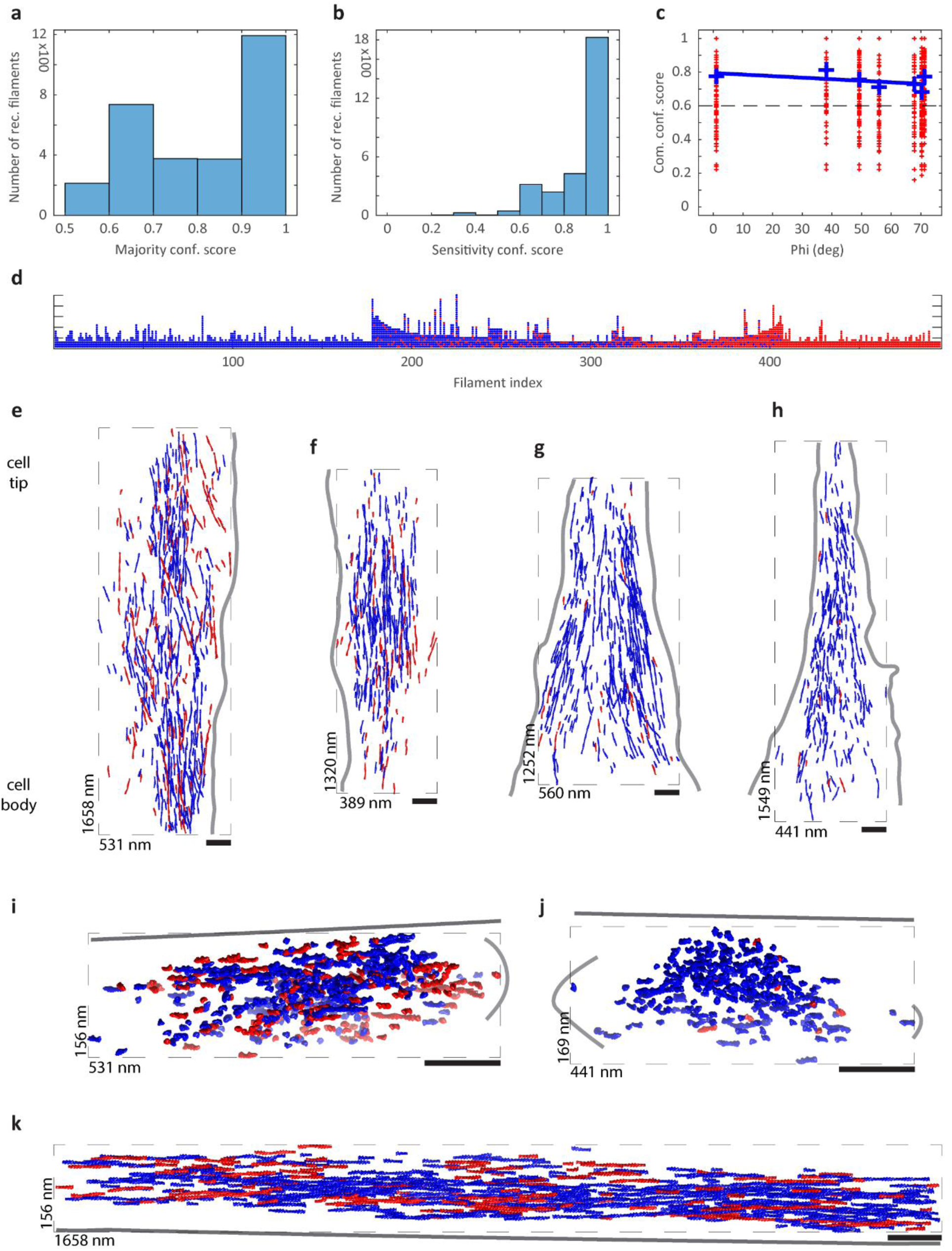
Architecture of actin bundles at FAs. (**a**) Histogram of the mcs values from 2893 reconstructed actin filaments, extracted from seven manually segmented cryo-tomograms acquired at FAs. (**b**) Histogram of the respective scs values. (**c**) Plot of the respective ccs values. Each red cross marks the ccs of one filament, and the scores are plotted as a function of the bundle orientation *φ*. The blue crosses indicate the average of all ccs values that are originating from the same actin bundle, and a regression line was fitted to these values (blue line). Only filaments reaching a ccs≥0.6 (dashed line) were accepted, therefore 2146 reconstructed actin filaments were used for subsequent bundle visualization and topology analysis. (**d**) Visualization of the resulting 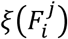 vectors of a bundle that was recorded at a proximal FA region. Segments, that are originating from the same filament are plotted as columns of circles. Blue segments are oriented with their plus-ends towards the cell tip, and red segments point in the direction of the cell body. In this bundle 495 filaments with an average length of 7.1 segments were analysed. Approximately 2/3 of the segments are directed towards the cell tip. (**e-h**) Top views on the architecture and polarity distribution of four actin bundles, recorded at proximal (**e**,**f**) and distal (**g**,**h**) FA regions. Blue actin filaments are oriented with their plus-ends towards the cell tip, and red filaments point in the direction of the cell body. Proximal FA bundles are characterized by a tendentially mixed polarity distribution, and distal FA bundles converge to a predominantly uniform polarity distribution. (**i**,**j**) Side views on the bundles shown in (**e**,**h**), respectively. (**i**) Filaments, which are directed towards the cell body (red filaments), are found with a higher probability at outer regions of proximal FA actin bundles. This relation is displayed in (**k**) as well, showing the long axis of bundle (**e**). In all visualizations (**e-k**), actin filaments were rendered from EMD-6179 (Galkin et al., 2015), the dashed rectangles are bounding boxes surrounding the actin bundles. Scale bars 100 nm. Additionally, positions of plasma membrane are suggested by light grey lines, and support planes by dark grey lines.

The segmentations can be conducted on contrast enhanced tomograms in order to facilitate a better detection of actin filaments. We do not assume the segmentations to be free of false positives, because they will be efficiently sorted out at a later stage. However, we assume that each filament or detected part is represented by a unique set of voxels, therefore filaments should not touch each other in the segmentations. The fact that a filament may appear divided into parts, for instance when it runs through low-contrast, dense or crowded regions, where it cannot be tracked unambiguously, has no detrimental effect on its polarity determination, but leads to an underestimation of the actual filament length.

The workflow of APT is shown in Fig. 1a. A detailed description of the consecutive steps can be found in the Method Details section.

### Validation of APT

The missing wedge affects the reconstruction of a tomogram in an anisotropic manner (Lucic et al., 2005). As a consequence, filaments running parallel to the tilt-axis are better resolved than filaments oriented orthogonal to the tilt-axis (Supplemental Fig. 1). It is fundamental to ensure that our projection approach for subtomogram averaging is not biased by the missing wedge. Additionally, the impact of SNR and polarity distribution on the output precision of APT should be verified.

For validation purposes we implemented a ground truth data generator (see Method Details), that creates modelled tomograms of actin bundles (volume 353 × 353 × 353 nm^3^ with a pixelsize of 3.44 Å), utilizing EMD-6179 (Galkin et al., 2015) as filament density, with three adjustable parameters: the angle *φ* of the bundle with the tilt-axis, the SNR of the tomogram (Forster et al., 2008; Pei et al., 2016), and the polarity ratio *ρ*_*r*_, namely the fraction of filaments in the bundle oriented in opposite direction. Each modelled bundle comprises on average 73 filaments, with a uniform length distribution of 86-282 nm and a uniform distance distribution of 10-37 nm between neighboring filaments.

In total we created three ground truth datasets with three different SNRs, that are 0.01, 0.001, and 0.0001. Each dataset consists of ten tomograms of different bundles, that are oriented in 10° steps from *φ* = 0° (filaments ∥ to the tilt-axis) to *φ* = 90° (filaments ⊥ to the tilt-axis). Additionally, within each set we varied the polarity ratio *ρ*_*r*_ in triplets, meaning that the first four tomograms (*φ* = 0°, 10°, 20°, 30°) were modelled with *ρ*_*r*_ = 0.1,0.3,0.5,0.1, and so forth, except the last tomogram (*φ* = 90°) was constructed with a polarity ratio of 0.5.

A top view on one of the modeled bundles (*φ* = 30°, *ρ*_*r*_ = 0.1) is shown in Fig. 1b (without noise, only for visualization purposes). A second example is depicted in Fig. 1c. This bundle is oriented 70° with respect to the tilt-axis and the polarity ratio is 0.3. Its side view is displayed in Fig. 1d, with the viewing axis oriented parallel to the filaments. A bandpass filtered slice of a filament, that was boxed out of one of the ground truth tomograms (*φ* = 0°, SNR = 0.001), is shown and can be compared to a filament parallel to the tilt-axis as well, but modelled with a SNR of 0.0001 (Fig. 1e and f, respectively).

We applied the APT workflow on each of the three ground truth datasets independently. Since the segmentation of the filaments is given by the ground truth data, we first established the segment coordinates (*Step I*), with an equidistant spacing of 11 nm, corresponding to ∼1000 segments per modelled bundle. Next, the subtomograms were extracted using a box size of 50 × 50 × 50 nm^3^ (*Step II*) and subsequently projected using a projection thickness of 11 nm (*Step III*). In the next step segments were prealigned (*Step IV*), followed by the 2D classification module (*Step V*). In Supplemental Fig. 4 the results of prealignment and 2D classification are shown for the modelled dataset with SNR = 0.0001.

**Figure 4.**
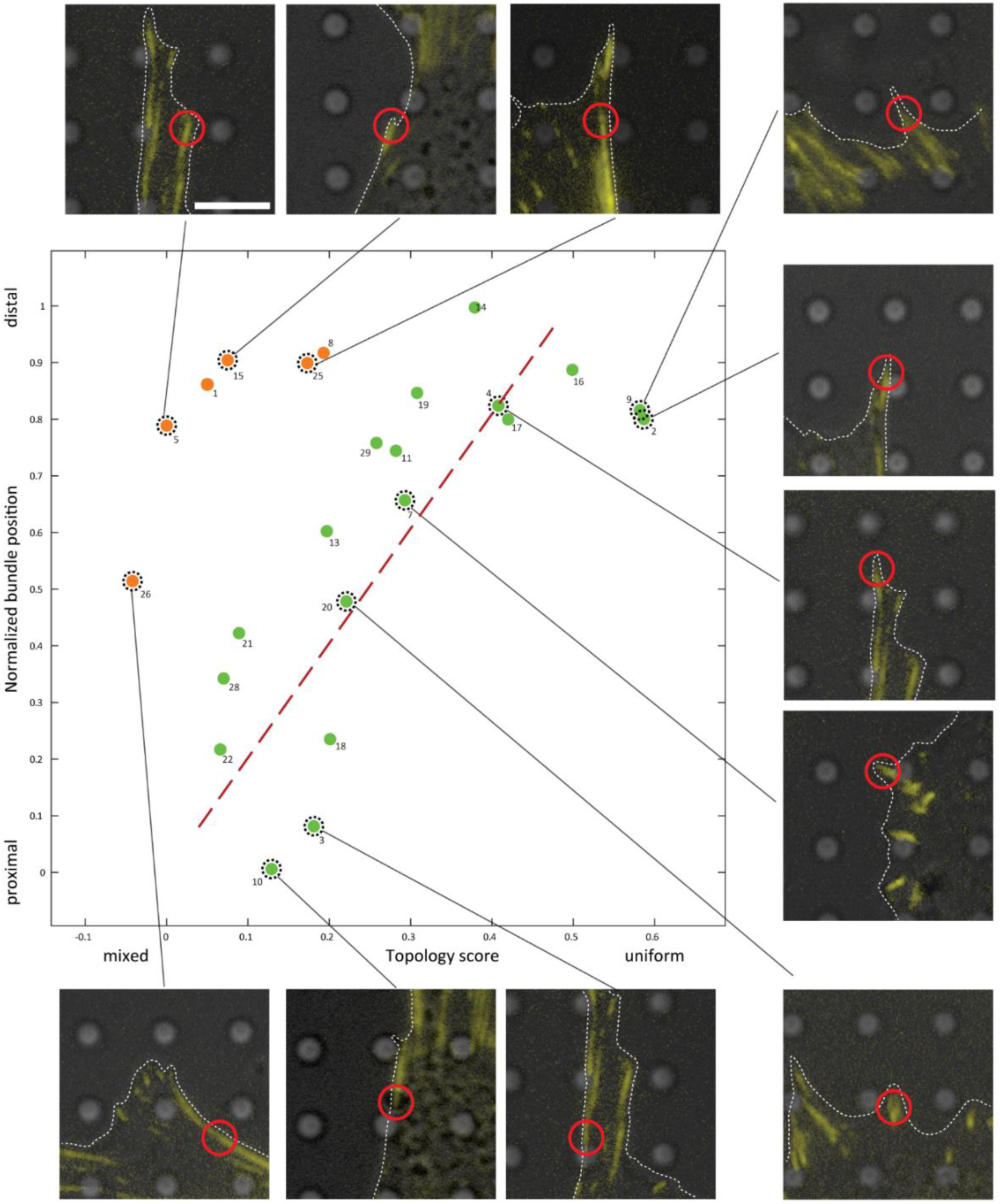
The distribution of actin polarity at FAs. Plot of topology score versus normalized bundle position along a FA. The green datapoints (fitted by the red dashed regression line) suggest a correlation between these two parameters. The polarity distribution along FAs transitions smoothly from mixed to uniform. However, in case that the distal region of a FA does not coincide with a pronounced cell protrusion, the polarity distribution exhibits a tendentially mixed character (orange datapoints). Some datapoints in the plot are accompanied by their fluorescence microscopy signal. The plasma membrane is outlined by white dashed lines, and the tomogram position is indicated by a red circle. Scale bar 5 µm.

Following this, we executed 3D reconstruction (*Step VI*) of the three actin filament averages from a similar number of particles (∼ 10’000 segments per average), distributed approximately uniform over the modelled bundles. As expected, the resolutions of the filament structures, drop with decreasing SNR (Supplemental Fig. 5).

**Figure 5.**
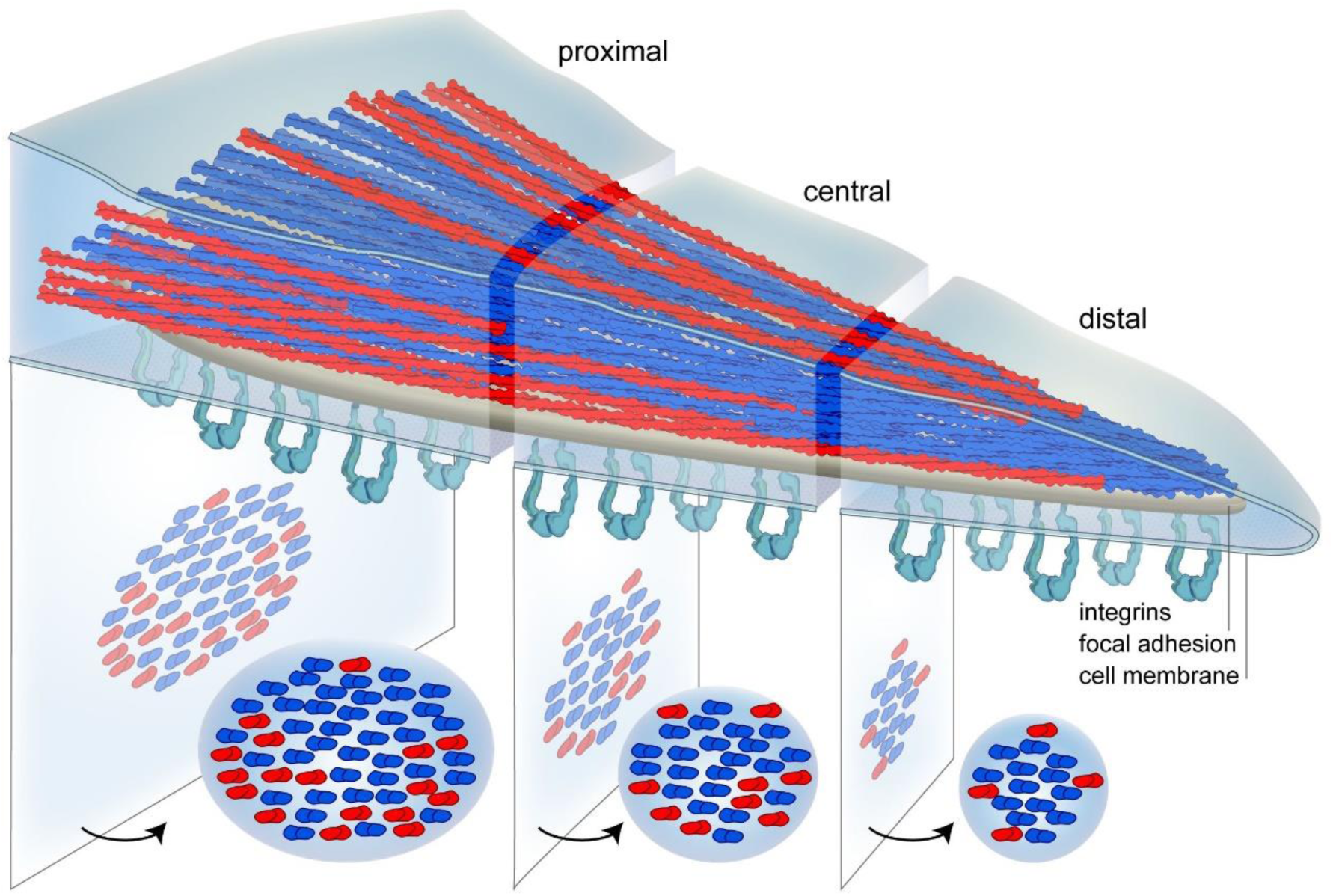
A model for actin polarity at FAs. Actin bundles within FAs are characterized by a mixed polarity distribution at proximal regions. However, there is a smooth transition towards a uniform polarity distribution at distal regions. Actin filaments, which are directed with their plus-ends towards the cell body (red), are arranged around a central core of filaments pointing towards the cell periphery (blue).

Subsequently, we performed polarity determination (*Step VII*) of all modelled bundles. All *ψ*^−1^ histograms (see Method Details) showed two distinct peaks (Fig. 1g), separated by 180°, indicating the opposite plus-end orientations of the segments, and the height of the peaks reproduce the respective polarity ratios. All restored bundles from all three SNR conditions showed satisfactory agreement with the ground truth bundles (Fig. 1h-i). Furthermore, we visualized the 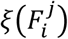 vectors (see Method Details) of all the recovered bundles (Fig. 1j). The position of incorrect assigned segments is randomly distributed and their number increases with decreasing SNR.

The results of the confidence analysis (*Step VIII*) are plotted in Fig. 1k. The ccs values (see Method Details) of bundles with SNR = 0.001 dropped slightly in comparison to bundles with SNR = 0.01, but show no trend to lower ccs values with increasing *φ* angle. We conclude, that the missing wedge induced anisotropic deterioration does not negatively influence the precision of APT for low SNR values, as found in cellular cryo-tomograms (Forster et al., 2008; Pei et al., 2016). However, for the markedly challenging SNR = 0.0001 the ccs values decrease with increasing *φ* angle.

Finally, we evaluated the total segment error (Fig. 1k), namely the fraction of segments with incorrect orientation determination compared to the ground truth. For low SNR conditions the total segment error is ≲10% for all modelled bundles, and increases to ∼20% for bundles modelled with SNR = 0.0001. However, if we evaluate the total filament error, that is the fraction of filaments with incorrect polarity determination compared to the ground truth, we find that the polarity of solely 5% of the filaments (34 out of 712 filaments) were determined incorrectly for the dataset modelled with SNR = 0.0001. This shows that APT is capable to reliably correct noise induced polarity errors on the filament level.

### Reconstruction of actin filaments at Fas

Here we applied APT on the actin cytoskeleton inside FAs, using a correlative fluorescence microscopy and cryo-ET approach (Patla et al., 2010; Sartori et al., 2007) (see Method Details). Mouse embryonic fibroblasts (MEFs), expressing vinculin-venus as a marker for FAs (Grashoff et al., 2010; Ringer et al., 2017), were cultured on electron microscopy grids with silicon-oxide support and imaged by fluorescence microscopy (Fig. 2a). Conspicuous FAs were identified and found again under the electron beam.

Due to the sheer size of FAs, tomograms cannot cover a complete adhesion site (∼2-4 µm in length), and due to electron dose sensitivity of the sample, only a single tomogram can be acquired per FA. Therefore, we recorded cryo-tomograms at a spectrum of positions, namely from proximal regions, where a stress fibre enters the FA, to distal regions, where they are oriented towards the plasma membrane. Seven cryo-tomograms of FAs (Fig. 2b), covering positions from proximal to distal, were selected in the first place, and the actin filaments were manually segmented (Fig. 2c).

Following this, we applied the APT workflow on this dataset. Firstly, we extracted a total of 43’400 segments with an equidistant spacing of 11 nm (*Step I*). The box size of the subtomograms was set to 50 × 50 × 50 nm^3^ (*Step II*), and the projection thickness to 11 nm (*Step III*). After prealignment and 2D classification (*Step IV-V*), we finally selected 20’585 segments (Supplemental Fig. 6a) for 3D reconstruction (*Step VI*).

The obtained in-situ actin filament structure was resolved to 18.4 Å, and allows to unambiguously detect the position of its plus-end (Supplemental Fig. 6b-d). This shows that the APT workflow is capable to produce sufficient resolution for subsequent mapping of the filament’s polarity distribution, although the dataset was relatively limited and the filaments originate from a crowded and dense cellular environment, with multiple possible binding partners and modulations of their helical symmetry.

In order to efficiently increase the size of the dataset, we used the previous seven manual segmentations to train a convolutional neural network with EMAN2 (Chen et al., 2017), capable of detecting actin filaments (see Method Details). Using this approach, we additionally created 31 segmentations of the actin cytoskeleton at FAs.

We then applied the APT workflow to this extended dataset. Here we extracted a total of 247’940 segments (with spacing of 11 nm), using a box size of 36 × 36 × 36 nm^3^ and a projection thickness of 11 nm (*Step I-III*). After prealignment and 2D classification (*Step IV-V*), we selected 72’973 segments (Supplemental Fig. 7, Fig. 2d) for 3D reconstruction (*Step VI*). Subsequently, we performed a 3D classification with RELION (Supplemental Fig. 8, Supplemental Fig. 9). The highest resolved class average with 13.5 Å is shown in Fig. 2e, together with a docking of EMD-6179 (Galkin et al., 2015) in Fig. 2f.

### Polarity distribution of actin bundles at Fas

Finally, we utilized APT to reconstruct actin networks in-situ, including the polarity of the filaments.

Therefore, we continued the APT workflow with the polarity determination module (*Step VII*), based on the in-situ actin filament reconstruction (Supplemental Fig. 6), previously obtained from seven manual segmentations of actin bundles at FAs. All *ψ*^−1^ histograms showed two distinct peaks, separated by 180°. We extracted the 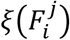 vectors of the resulting 2893 filaments, and used APT to calculate the confidence scores (*Step VIII*), plotted in Fig. 3a-c. The minimum reliable ccs was set to 0.6 (Fig. 3c), thereby leaving a total amount of 2146 actin filaments with determined polarity. The total length of the filaments is ∼149 µm, with an average length of ∼70 nm, and a maximum length of ∼350 nm (Supplemental Fig. 10a,b).

In Fig. 3d the resulting 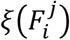 vectors of one of the bundles are displayed. Here we found that 62% of the segments are pointing with their plus-ends towards the cell tip (blue segments), and 38% in opposite direction (red segments). The actin polarity distribution of this actin bundle is visualized in Fig. 3e. The blue filaments are pointing with their plus-ends towards the cell tip, and the red filaments in opposite direction.

The two bundles shown in Fig. 3e-f were acquired at proximal regions of FAs, whereas the two bundles shown in Fig. 3g-h were acquired at distal regions. We applied APT’s topology module (*Step IX*), and sorted the bundles with increasing topology score (Supplemental Fig. 10c) from left (*τ*=0.25) to right (*τ*=0.87), which reflects their transition from a tendentially mixed polarity distribution to a predominantly uniform polarity distribution. In Fig. 3e, 65% of the filaments are directed towards the cell tip, whereas in Fig. 3h this fraction is increased to 96%.

The bundles in Fig. 3e and Fig. 3h are shown from the side in Fig. 3i and Fig. 3j. Furthermore, the long axis of the bundle in Fig. 3e is shown in Fig. 3k. The data suggests that filaments, which are directed towards the cell tip, forming the core of the FA actin bundle, and the reversely directed filaments are organized around this core. The positive topology scores of bundles with mixed polarity distributions are a further indication for this architectural principle. If the reversed actin filaments would be diffused into the bundles, the topology score would approach negative values. In addition, we analysed the polarity distribution in layers parallel to the support, finding that reversed filaments are enriched on top and bottom of the mixed polarity bundles (Supplemental Fig. 10d).

The number of filaments within neighbourhood spheres 𝔎_*ϵ*_(*x*_*k*_), which were evaluated during topology analysis (*Step IX*) with a radius of *ϵ*=40 nm, has its maximum at 6 neighbours (Supplemental Fig. 10e). This indicates that actin filaments in FA bundles resemble a hexagonal packing with a mean filament distance of 28 nm (Supplemental Fig. 10f).

Finally, we completed the APT workflow (*Step VII-IX*) for the extended dataset with previously obtained 31 automatic segmentations (Fig. 2d-f, Supplemental Fig. 7-9). Seven tomograms were excluded, because they contained ill-defined or multiple bundles with similar orientation, so that the *ψ*^−1^ histograms did not exhibit two clearly defined peaks. As before, we set the minimal reliable ccs to 0.6, thereby leaving a total amount of 6152 actin filaments with determined polarity. The total length of the analyzed filaments was increased to ∼388 µm. In Fig. 4 we plotted the topology score *τ* of these 24 bundles versus their normalized position along a FA (see Method Details). Strikingly, a pronounced uniform polarity distribution (high *τ* values) can be found in distal regions of FAs. For most of the analyzed bundles (Fig. 4, green dots) the plot suggests a correlation between topology score and normalized FA position. In other words, within a FA, from proximal to distal regions, the polarity distribution of the actin filaments transforms smoothly from mixed to uniform actin polarity organization. Interestingly, this correlation does not hold true, when the FA is not positioned towards a pronounced cell protrusion (Fig. 4, orange dots).

## Discussion

Resolving the polarity of actin filaments inside cells, would provide detailed insights into cytoskeletal processes and their functional organization. Therefore, we implemented APT, a set of MATLAB scripts and RELION procedures that enables the reconstruction of actin filaments and the assessment of their polarity. This general approach can be applied to any cryo-tomograms, in which actin filaments are observed. Moreover, the usage of APT to reconstruct in-situ actin filament structures, opens new avenues to explore complex actin networks and even structural interactions of actin and actin binding proteins.

The basic concept of APT is to dissect an actin filament into multiple segments, and subsequently measure their polarity with single particle techniques. Thereby we create a set of statistically independent estimations of the filament’s polarity. Based on majority, the final polarity of the filament is determined. We validated this approach and found that the total filament error is below 5%, even for very challenging and low SNR = 0.0001.

A previous approach to decipher actin polarity from tomographic data necessitated chemical extraction and negative staining of the cytoskeleton (Narita et al., 2012), in order to increase the SNR of the tomograms. However, the challenge of resolving actin filament structure and polarity in-situ requires a tomographic data set from intact cells (Beck and Baumeister, 2016; Lucic et al., 2005), which involves vitrification of the sample, and therefore the method needs to successfully handle low SNR (Forster et al., 2008; Pei et al., 2016).

As an example, we applied APT on actin bundles found at FAs. These multi-layered assemblies, around 200-350 nm in thickness, connect integrin receptors, which anchor the plasma membrane to the extracellular matrix, via force transduction and actin regulation layers, with the actin cytoskeleton (Kanchanawong et al., 2010; Patla et al., 2010).

Actin bundles at FAs (Fig. 3e-k) resemble a hexagonal packing of the filaments with an average distance of ∼28 nm (Supplemental Fig. 10e,f). The majority of the filaments is directed towards the cell tip (Fig. 3e-h), and form the core of the bundles, whereas filaments with reversed direction organize around this core (Fig. 3i-k, Supplemental Fig. 10d).

Ultimately, the polarity distribution depends on the position of the bundle along the FA (Fig. 4). The more distal the position, the more uniform the polarity distribution, while mixed polarity can be observed mainly in the proximal regions (Fig. 5). Stress fibers are contractile actin assemblies (Malik-Garbi et al., 2019), enriched with actomyosin interactions, which favor a mixed polarity distribution of actin filaments. Proximal regions of FAs anchor to stress fibers (Burridge and Guilluy, 2016), therefore exhibiting a mixed polarity actin bundle. However, at distal regions of FAs, more vectoral or even protruding forces may play a role that are resembled in a more uniform polarity distribution.

Technical developments in sample preparation allowed cryo-ET to provide fundamental insights into cellular assemblies and processes in-situ (Mahamid et al., 2016; Marko et al., 2007; Schaffer et al., 2019). However, it also requires the development of novel tools to analyse the data (Chen et al., 2017; Himes and Zhang, 2018; Martinez-Sanchez et al., 2020; Song et al., 2019). APT complements these tools and allows to analyse actin filaments in physiological relevant processes, which would provide a better understanding of actin networks, such as the actin cortex (Chugh et al., 2017), stereocilia organization (Metlagel et al., 2019) or remodelling of actin during phagocytosis (Gerisch, 2011).

## Method Details

### Implementation and code availability

All functionality of APT was implemented as MATLAB (MathWorks, Natick, USA) scripts and functions. In order to interface with RELION (Scheres, 2012) (tested with version 3.0.4), respective functions to write and read starfiles were programmed.

The software is available at https://www.placeholder.com. It is equipped with a tutorial that explains APT step-by-step based on modelled tomograms of actin bundles.

### Step I: Coordinates

The input data is a set of *N*_*T*_ cryo-electron tomograms *T*_*i*_ (*i* = 1, …, *N*_*T*_) and for each *T*_*i*_ there is an associated segmentation *S*_*i*_ of the actin filaments. Each *S*_*i*_ define a set of filaments 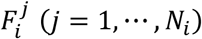), with *N*_*i*_ being the number of detected filaments in *T*_*i*_. Firstly, APT utilizes the *S*_*i*_ to construct for each 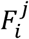 a set of 3D coordinates 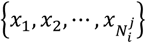, that are evenly spaced along 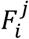, and will be the sampling points for polarity determination. Here 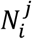 indicates the number of sampling points per filament, which we term segments.

### Step II: Subtomograms

Next, APT pools all these 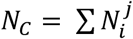 segments *x*_*k*_ (*k* = 1, …, *N*_*C*_), stores the index relation 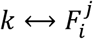 for polarity determination at a later stage, and extracts *N*_*C*_ subtomograms centred around *x*_*k*_ from a ctf-corrected version of the dataset *T*_*i*_.

### Step III: Projections

Subsequently, APT masks and then projects the subtomograms in z-direction. The applied mask diminishes the influence of neighbouring filaments in z-direction. We term the height of the mask as projection thickness parameter.

### Step IV: Prealignment

This is the first of three steps performed in RELION. First, a starfile was created only passing the name of each projected subtomogram (`rlnImageName`) and its originating tomogram (`rlnMicrographName`) to RELION. Next, the projected subtomograms were normalized with `relion_preprocess`. Please note that the index relation 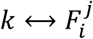 was kept invisible for RELION in all the following steps (for example no metadata label `rlnHelicalTubeID` was provided (He and Scheres, 2017)). Furthermore, CTF correction in RELION jobs was switched off, because the tomograms were CTF corrected by phase-flipping prior to subtomogram extraction.

For prealignment of the segments a 2D classification with the prepared starfile as input was performed. The purpose of this step is to align the (central) actin filament density parallel with the x-axis, which is the RELION convention for helical reconstruction (He and Scheres, 2017). Therefore, we created a set of projection images (the template library) from an actin filament structure, which was oriented along the x-axis, rotated in 3° increments and subsequently projected. The template library was used as initial references for prealignment (Supplemental Fig. 4a).

Since the resolution of the actin filament averages was determined by Fourier shell correlation between the averages and EMD-6179 (Galkin et al., 2015) as an external reference (Supplemental Fig. 5, Supplemental Fig. 6d, and Supplemental Fig. 9), we were reluctant to use the same structure for the construction of the template library. For that purpose, we used EMD-10737.

### Step V: 2D Classification

Subsequently we performed a second 2D classification in RELION, passing the psi angles and translations found in the previous step as priors (`rlnAnglePsiPrior`, `rlnOriginXPrior`, `rlnOriginYPrior`). In contrast to the prealignment step, it is crucial that this 2D classification is unsupervised, in order to extract the structural heterogeneity in the data unbiasedly (Supplemental Fig. 4c-f).

The psi search was conducted local around the psi prior. Therefore, it was possible to apply a second mask (the first mask was applied during the projection step in z-direction), which allowed to further diminish the influence of neighbouring filaments in y-direction.

This second 2D classification is capable of producing high quality class averages of filamentous actin from inside intact cells (Supplemental Fig. 6a, Supplemental Fig. 7), and allows to sort out false positive actin detections and low-quality segments (Bharat and Scheres, 2016). Based on this classification we selected the segments for subsequent 3D reconstruction.

### Step VI: 3D Reconstruction

For actin filament averaging a 3D refine job in RELION was performed with helical reconstruction switched on (He and Scheres, 2017). As helical rise 27.6 Å and as helical twist 166.7° was used (Galkin et al., 2015). In the input starfile, the refined psi angle and translations found in the previous step were passed as priors plus for each segment a random rot angle as prior was added and the tilt prior was set to 90° (He and Scheres, 2017) (`rlnAngleRotPrior`, `rlnAngleTiltPrior`). As initial template the same structure as for construction of the template library was used, low-pass filtered to 30 Å (Supplemental Fig. 6d). Subsequently, APT combines all transformations, and as a result we have the forward transformation Ω_*k*_, that describes how filament 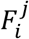 has to be transformed at sampling point *x*_*k*_ in order to align with the filament average (Supplemental Fig. 3a).

### Step VII: Polarity

To resolve actin polarity, it is crucial that the filament average reaches a resolution better than 20 Å. Once this has been achieved, APT calculates the inverse transformation 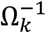, that describes how filament 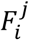 is oriented at sampling point *x*_*k*_ with respect to the filament average (Supplemental Fig. 3a). Particularly filament polarity is encoded in the inverted psi angle 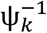, which indicates the position of the plus-end of 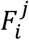 relative to the y-axis of *T*_*i*_.

Next, APT uses the index relation 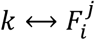 to extract all inverted psi angles 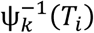, that are originating from the same *T*_*i*_. If *T*_*i*_ contains a filament bundle with mixed polarity, then the histogram of 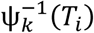 shows two distinct peaks, that are separated by 180°. The height of each peak indicates how many segments were aligned with their plus-ends in the respective direction, and allows an estimation of the polarity ratio within the bundle, while the peak width decreases with increasing parallelism of the filaments. Obviously, if a bundle exhibits a uniform polarity distribution, then the histogram of 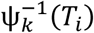 shows a single peak.

Subsequently, APT assigns to each segment a direction label. Per default, segments included in the right peak are labelled with 0, segments included in the left peak are labelled with 1. Therefore, each filament can be represented as a vector 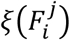 of labelled segments.

The centrepiece of APT is that the elements of 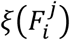 are computed statistically independent. Segments, whose directionality was detected incorrectly, do not prohibit the correct polarity determination of a filament as a whole, as long as the majority of segments building that filament were determined accurately. Consequently, the final polarity label 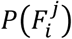 of filament 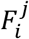 is defined as follows:

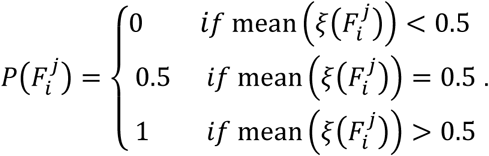

If 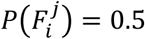 the polarity of that filament is maximally uncertain.

### Step VIII: Confidence

We suggest that the closer mean 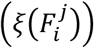 is to 0 or to 1, respectively, the higher the probability, that 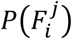 was determined correctly. We use this relation to define a confidence score for each filament, termed majority confidence score (mcs), as follows:

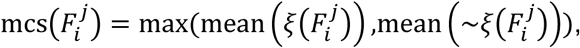

with 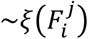 the logical complement of 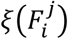. The values of mcs range between 0.5 (maximum uncertainty) to 1 (minimal uncertainty), and are the fraction of segments per filament that are pointing in the majority direction. The mcs measures the consistency of the segment’s directionality determination along a filament.

Additionally, we define a second confidence score, termed sensitivity confidence score (scs). Here the reconstruction of the filament average is repeated with a reversed direction of the template. Subsequently, APT initializes for each 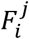 a vector 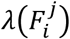, having the same number of elements than 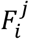 segments. Those segments that reverse their plus-end orientation as well, are orientational sensitive and APT sets the respective elements of 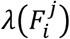 to 1. On the other hand, if a segment shows no logical behaviour upon change in template direction, the respective element of 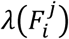 is set to 0. Then we define scs as follows:

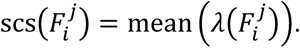

The values of scs range between 0 (no segment of 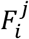 is orientational sensitive) and 1 (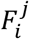 is built exclusively from orientational sensitive segments). Comparable to the mcs, the scs is an alternative to measure the self-consistency of the segment’s directionality determination. Finally, we define the combined confidence score (ccs) of filament 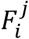 as follows:

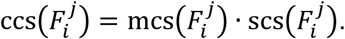

### Step IX: Topology

In this module APT performs a 3D neighbourhood analysis of the actin filaments in order to characterize their polarity distribution in the observed bundles. Therefore, centred around each segment *x*_*k*_, APT constructs a sphere 𝔎_*ϵ*_(*x*_*k*_) with radius *ϵ*, defining the neighbourhood of *x*_*k*_. Then, APT extracts number, distance and polarity of all filaments passing through 𝔎_*ϵ*_(*x*_*k*_). For instance, let the polarity of the segment *x*_*k*_ be 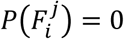, and imagine 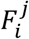 is surrounded by four filaments within 𝔎_*ϵ*_(*x*_*k*_), of which one displays same polarity and three reversed polarity (Supplemental Fig. 3b). Then, the position of the probed filament is located in a neighbourhood that exhibits 75% mixed polarity and 25% uniform polarity, respectively. This calculation is performed by APT for all 𝔎_*ϵ*_(*x*_*k*_), and subsequently it sums the degrees of uniform polarity *P*_∥_ and mixed polarity *P*_∦_ of all neighbourhoods in a given actin bundle. Finally, the topology score *τ* of an actin bundle is defined as follows:

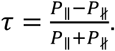

The topology score *τ* ranges from −1 to +1. The more the filaments are organized in a uniform polarity configuration, the value of *τ* will be closer to +1. If the bundle shows a phase separated topology, so that filaments with opposite polarities adjoin at phase boundaries mainly, *τ* will be reduced accordingly. However, the more the bundle favours close proximity between filaments with opposite polarities, the more *τ* will approach −1.

### Ground truth data generator

In the first step of modelling a cryo-tomogram of an actin bundle, filament x-z-coordinates were calculated based on random close packing (Desmond and Weeks, 2009), with a uniform distance distribution in the range between 10-37 nm. As a consequence of the modelled x-z-dimension of the bundles of 282 x 150 nm, on average 73 filaments could be packed in each bundle. Next, the position and length of each filament in y-direction was randomly chosen from a uniform distribution between 86-282 nm.

According to these coordinates, actin filaments based on EMD-6179 (Galkin et al., 2015), were pasted in an empty volume of 353 × 353 × 353 nm^3^, however, a random fraction of *ρ*_*r*_ filaments was pasted in the volume after 180° rotation around the x-axis (Fig. 1b-d, red coloured filaments).

After that the modelled bundle was rotated *φ* degrees around the z-axis, then Gaussian noise was added to adjust the targeted SNR, and finally the missing wedge of a tilt-series with tilt-range from −60° to +60° was applied in Fourier space.

### Cell culture

MEFs stably expressing vinculin-venus were cultured in Dulbecco’s Modified Eagle’s Medium (Sigma-Aldrich, D5671), supplemented with 10% (v/v) fetal bovine serum (Sigma-Aldrich, G7524), 2 mM L-glutamine (Sigma-Aldrich, G7513) and 100 µg/ml penicillin-streptomycin (Sigma-Aldrich, P0781), at 37° C and 5% CO_2_. The MEFs were applied onto glow-discharged EM grids with a silicon-oxide support film (R1/4, Au mesh; Quantifoil, Jena, Germany). After incubation, the cells were fixed in a 4% paraformaldehyde solution (Sigma-Aldrich, 16005), and washed 3 times with 1x PBS (Fisher Scientific, BP399-1).

### Fluorescence microscopy

The EM grids were transferred to a 35×14 mm glass-bottom cell culture dish (MatTek, P35G-0-14-C) for fluorescence microscopy analysis of the FAs. Adherent MEFs were imaged using an automated inverted microscope (DMI4000 B, Leica Microsystems, Wetzlar, Germany) equipped with a fluorescence lamp and a monochromatic digital camera (DFC365 FX, Leica Microsystems). Overview images of the EM grids were acquired using a 10x dry objective (HCX PL Fluotar 10x/0.30, Leica Microsystems). All images of individual cells (Fig. 2a) were acquired in phase contrast and fluorescence mode using a 63x oil objective (HCX PL APO 63x/1.40-0.6, Leica Microsystems).

### Cryo-electron tomography

A drop of 4 µl BSA-coated 10 nm fiducial gold markers (Aurion, Wageningen, Netherlands) was applied on the EM grids before plunging them into liquid nitrogen cooled ethane.

A Titan Krios transmission electron microscope (Thermo Fisher Scientific, Waltham, USA) equipped with a Quantum energy filter and a K2-Summit direct electron detector (Gatan, Pleasanton, USA) was used for cryo-EM data acquisition. The microscope was operated at 300 keV in zero-loss mode with the energy filter slit width set to 20 eV.

The position of the cells and FAs were identified by overlaying the fluorescence signal to low magnification EM images of the grids. At tomography positions image stacks were recorded at each tilt angle in super-resolution mode with an electron flux of ∼8 electrons per pixel per second using SerialEM (Mastronarde, 2005). All tomograms were acquired at a magnification of 42’000x, and a dose-fractionated frame rate of 6 frames per 1.2 s. The tilt-series covered an angular range of −60° to +60°, and were recorded with tilt increments of 2° and a defocus of −4 μm. The accumulated electron dose did not exceed ∼75 e^-^/Å^2^.

All image stacks were down-sampled and motion corrected using MotionCorr (Li et al., 2013), resulting in a final pixel size of 3.44 Å. Next, CTF correction of the tilt-series was applied (Eibauer et al., 2012). Overview tomograms (Fig. 2b) for actin filament segmentation (Fig. 2c) and subtomograms of actin segments were reconstructed by weighed back-projection using the TOM Toolbox (Nickell et al., 2005).

### Normalized bundle position of FA

The tomographic position within a FA was determined by overlaying the light microscopy images of the individual cells with cryo-EM images. First, phase contrast and fluorescence images were overlaid, hereby enabling the identification of the cell membrane and enclosing grid holes. Second, low magnification EM images (4’800x) of the FA tomography position were overlaid such, that the cell membrane and grid holes coincided with the light microscopy images. Third, the zero-degree tilt-series image (42’000x) was aligned with the low magnification EM image such, that the cell membrane shape matched. By increasing the opacity of the EM images, we displayed the FA fluorescence signal on the tomography position.

We defined the center of the tomogram as the tomography position FA_T_. In order to determine the relative position of FA_T_ within the entire FA, we defined the proximal end (closest to cell body) and distal end (closest to cell periphery) fluorescence microscopy signal pixels of the FA as FA_P_ and FA_D_, respectively. The distance *d* between FA_T_ and FA_D_ and the length *l* of the FA (distance between FA_P_ and FA_D_) was measured (the pixel size of the fluorescence microscopy images was 91 nm). Finally, the normalized bundle position of the FA (plotted on the y-axis in Fig. 4) was calculated as 1 − *d*/*l*.

### Automatic segmentation

We developed a script in MATLAB to automatize the convolutional neural network (CNN) segmentation with EMAN2 (Chen et al., 2019; Chen et al., 2017). The method requires three image stacks with fixed x-y-dimension of 64 x 64 pixels as input files. Firstly, the positive training stack contains 2D images of densities targeted for segmentation. These images are slices, boxed out of cryo-tomograms. In our case they contain actin filament densities. Secondly, an image stack that provides for each image in the positive training stack an associated segmentation mask. Thirdly, the negative training stack, which contains 2D images of densities not targeted for segmentation, for example gold particles used as fiducial markers, backprojection rays, tomogram edges, plasma membrane, and regions without actin filaments in general.

As a starting point we created seven manual segmentations of actin filaments in cryo-tomograms of FA actin bundles (Fig. 2c) with AMIRA (Thermo Fisher Scientific, Waltham, USA). The segmentations were skeletonized, then the filaments were dissected in segments, and around each 3D coordinate of a segment one positive training image was extracted from the tomogram at respective z-position. For each manual segmentation one positive training stack was created, limited to 3’000 images.

In order to create the associated segmentation masks, we extended the skeletonized filaments to the diameter of an actin filament. Subsequently, the segmentation masks were extracted from the extended segmentation at the same coordinates as the positive training images. For each manual segmentation one stack with segmentation masks was created, with the same dimension as the associated positive training stack.

Negative training images were extracted at random positions from the tomograms, but the volume of the actin bundle was excluded. For each manual segmentation one negative training stack was created, limited to 3’000 images as well.

Based on these image stacks, we trained one CNN for each manually segmented tomogram, using default EMAN2 data augmentation, network and training parameters. In the next step, the resulting seven CNNs were applied to each of the 31 tomograms in the extended dataset, thereby creating a total amount of 217 segmentations. Finally, the segmentations belonging to the same tomogram were averaged and post-processed in UCSF Chimera (Pettersen et al., 2004) with the hide dust command.

### Visualization

All isosurface visualizations of actin filament structures and actin bundles were rendered with UCSF Chimera or AMIRA.

For the visualizations of actin bundles in Fig. 3e-k we used EMD-6179 (Galkin et al., 2015) to represent the filaments. Therefore, we used a b-spline registration algorithm to bend the filaments in order to match their 3D shape defined by the segment coordinates (Rueckert et al., 1999).

## Data availability

Actin filament averages will be deposited in the Electron Microscopy Data Bank under accession codes EMD-abcd_1_ for the actin filament structure obtained from the manual segmented dataset (Supplemental Fig. 6), and EMD-abcd_2_, EMD-abcd_3_, and EMD-abcd_4_ for the actin filament class averages (Supplemental Fig. 8) obtained from the extended dataset.

## Acknowledgements

This work was funded by grants from the Swiss National Science Foundation (SNSF 31003A_179418), ERC-Syg and the Mäxi Foundation to O.M. We thank Carsten Grashoff for the vinculin labelled MEF cells. We thank the Center for Microscopy and Image Analysis at the University of Zurich.

## Author contributions

B. M. prepared samples and recorded data. M. E. developed and implemented the method. B. M., S. S., W. C., and M. T. wrote and tested code. B. M. and M. E. analyzed data. O. M. and M. E. conceived the research and wrote the manuscript with input from all authors.

## References

Beck, M., and Baumeister, W. (2016). Cryo-Electron Tomography: Can it Reveal the Molecular Sociology of Cells in Atomic Detail? Trends Cell Biol 26, 825–837.

Bharat, T.A.M., and Scheres, S.H.W. (2016). Resolving macromolecular structures from electron cryo-tomography data using subtomogram averaging in RELION. Nat Protoc 11, 9–20.

Burridge, K., and Guilluy, C. (2016). Focal adhesions, stress fibers and mechanical tension. Exp Cell Res 343, 14–20.

Chen, M., Bell, J.M., Shi, X., Sun, S.Y., Wang, Z., and Ludtke, S.J. (2019). A complete data processing workflow for cryo-ET and subtomogram averaging. Nat Methods 16, 1161–1168.

Chen, M., Dai, W., Sun, S.Y., Jonasch, D., He, C.Y., Schmid, M.F., Chiu, W., and Ludtke, S.J. (2017). Convolutional neural networks for automated annotation of cellular cryo-electron tomograms. Nat Methods 14, 983–985.

Chugh, P., Clark, A.G., Smith, M.B., Cassani, D.A.D., Dierkes, K., Ragab, A., Roux, P.P., Charras, G., Salbreux, G., and Paluch, E.K. (2017). Actin cortex architecture regulates cell surface tension. Nat Cell Biol 19, 689–697.

Desmond, K.W., and Weeks, E.R. (2009). Random close packing of disks and spheres in confined geometries. Phys Rev E 80.

Egelman, E.H., Francis, N., and DeRosier, D.J. (1982). F-actin is a helix with a random variable twist. Nature 298, 131–135.

Eibauer, M., Hoffmann, C., Plitzko, J.M., Baumeister, W., Nickell, S., and Engelhardt, H. (2012). Unraveling the structure of membrane proteins in situ by transfer function corrected cryo-electron tomography. Journal of Structural Biology 180, 488–496.

Elad, N., Volberg, T., Patla, I., Hirschfeld-Warneken, V., Grashoff, C., Spatz, J.P., Fassler, R., Geiger, B., and Medalia, O. (2013). The role of integrin-linked kinase in the molecular architecture of focal adhesions. J Cell Sci 126, 4099–4107.

Forster, F., Medalia, O., Zauberman, N., Baumeister, W., and Fass, D. (2005). Retrovirus envelope protein complex structure in situ studied by cryo-electron tomography. Proc Natl Acad Sci U S A 102, 4729–4734.

Forster, F., Pruggnaller, S., Seybert, A., and Frangakis, A.S. (2008). Classification of cryo-electron sub-tomograms using constrained correlation. J Struct Biol 161, 276–286.

Galkin, V.E., Orlova, A., Vos, M.R., Schroder, G.F., and Egelman, E.H. (2015). Near-atomic resolution for one state of F-actin. Structure 23, 173–182.

Geiger, B., Spatz, J.P., and Bershadsky, A.D. (2009). Environmental sensing through focal adhesions. Nat Rev Mol Cell Biol 10, 21–33.

Gerisch, G. (2011). Actin switches in phagocytosis. Commun Integr Biol 4, 344–345.

Grashoff, C., Hoffman, B.D., Brenner, M.D., Zhou, R., Parsons, M., Yang, M.T., McLean, M.A., Sligar, S.G., Chen, C.S., Ha, T., et al. (2010). Measuring mechanical tension across vinculin reveals regulation of focal adhesion dynamics. Nature 466, 263–266.

He, S., and Scheres, S.H.W. (2017). Helical reconstruction in RELION. J Struct Biol 198, 163–176.

Hervas-Raluy, S., Garcia-Aznar, J.M., and Gomez-Benito, M.J. (2019). Modelling actin polymerization: the effect on confined cell migration. Biomech Model Mechanobiol 18, 1177–1187.

Himes, B.A., and Zhang, P. (2018). emClarity: software for high-resolution cryo-electron tomography and subtomogram averaging. Nat Methods 15, 955–961.

Holmes, K.C., Popp, D., Gebhard, W., and Kabsch, W. (1990). Atomic model of the actin filament. Nature 347, 44–49.

Jasnin, M., Beck, F., Ecke, M., Fukuda, Y., Martinez-Sanchez, A., Baumeister, W., and Gerisch, G. (2019). The Architecture of Traveling Actin Waves Revealed by Cryo-Electron Tomography. Structure 27, 1211–1223 e1215.

Kanchanawong, P., Shtengel, G., Pasapera, A.M., Ramko, E.B., Davidson, M.W., Hess, H.F., and Waterman, C.M. (2010). Nanoscale architecture of integrin-based cell adhesions. Nature 468, 580–584.

Legate, K.R., Takahashi, S., Bonakdar, N., Fabry, B., Boettiger, D., Zent, R., and Fassler, R. (2011). Integrin adhesion and force coupling are independently regulated by localized PtdIns(4,5)2 synthesis. EMBO J 30, 4539–4553.

Li, X.M., Mooney, P., Zheng, S., Booth, C.R., Braunfeld, M.B., Gubbens, S., Agard, D.A., and Cheng, Y.F. (2013). Electron counting and beam-induced motion correction enable near-atomic-resolution single-particle cryo-EM. Nature Methods 10, 584-+.

Lucic, V., Forster, F., and Baumeister, W. (2005). Structural studies by electron tomography: from cells to molecules. Annu Rev Biochem 74, 833–865.

Mahamid, J., Pfeffer, S., Schaffer, M., Villa, E., Danev, R., Cuellar, L.K., Forster, F., Hyman, A.A., Plitzko, J.M., and Baumeister, W. (2016). Visualizing the molecular sociology at the HeLa cell nuclear periphery. Science 351, 969–972.

Malik-Garbi, M., Ierushalmi, N., Jansen, S., Abu-Shah, E., Goode, B.L., Mogilner, A., and Keren, K. (2019). Scaling behaviour in steady-state contracting actomyosin networks. Nat Phys 15, 509–516.

Marko, M., Hsieh, C., Schalek, R., Frank, J., and Mannella, C. (2007). Focused-ion-beam thinning of frozen-hydrated biological specimens for cryo-electron microscopy. Nat Methods 4, 215–217.

Martinez-Sanchez, A., Kochovski, Z., Laugks, U., Meyer Zum Alten Borgloh, J., Chakraborty, S., Pfeffer, S., Baumeister, W., and Lucic, V. (2020). Template-free detection and classification of membrane-bound complexes in cryo-electron tomograms. Nat Methods.

Mastronarde, D.N. (2005). Automated electron microscope tomography using robust prediction of specimen movements. Journal of Structural Biology 152, 36–51.

Medalia, O., Weber, I., Frangakis, A.S., Nicastro, D., Gerisch, G., and Baumeister, W. (2002). Macromolecular architecture in eukaryotic cells visualized by cryoelectron tomography. Science 298, 1209–1213.

Merino, F., Pospich, S., Funk, J., Wagner, T., Kullmer, F., Arndt, H.D., Bieling, P., and Raunser, S. (2018). Structural transitions of F-actin upon ATP hydrolysis at near-atomic resolution revealed by cryo-EM. Nat Struct Mol Biol 25, 528–537.

Metlagel, Z., Krey, J.F., Song, J., Swift, M.F., Tivol, W.J., Dumont, R.A., Thai, J., Chang, A., Seifikar, H., Volkmann, N., et al. (2019). Electron cryo-tomography of vestibular hair-cell stereocilia. J Struct Biol 206, 149–155.

Narita, A., Mueller, J., Urban, E., Vinzenz, M., Small, J.V., and Maeda, Y. (2012). Direct determination of actin polarity in the cell. J Mol Biol 419, 359–368.

Nickell, S., Forster, F., Linaroudis, A., Net, W.D., Beck, F., Hegerl, R., Baumeister, W., and Plitzko, J.M. (2005). TOM software toolbox: acquisition and analysis for electron tomography. J Struct Biol 149, 227–234.

Nogales, E. (2016). The development of cryo-EM into a mainstream structural biology technique. Nat Methods 13, 24–27.

Patla, I., Volberg, T., Elad, N., Hirschfeld-Warneken, V., Grashoff, C., Fassler, R., Spatz, J.P., Geiger, B., and Medalia, O. (2010). Dissecting the molecular architecture of integrin adhesion sites by cryo-electron tomography. Nat Cell Biol 12, 909–915.

Pei, L., Xu, M., Frazier, Z., and Alber, F. (2016). Simulating cryo electron tomograms of crowded cell cytoplasm for assessment of automated particle picking. BMC Bioinformatics 17, 405.

Pettersen, E.F., Goddard, T.D., Huang, C.C., Couch, G.S., Greenblatt, D.M., Meng, E.C., and Ferrin, T.E. (2004). UCSF chimera - A visualization system for exploratory research and analysis. J Comput Chem 25, 1605–1612.

Pollard, T.D., and Borisy, G.G. (2003). Cellular motility driven by assembly and disassembly of actin filaments. Cell 112, 453–465.

Punjani, A., Rubinstein, J.L., Fleet, D.J., and Brubaker, M.A. (2017). cryoSPARC: algorithms for rapid unsupervised cryo-EM structure determination. Nat Methods 14, 290–296.

Ringer, P., Weissl, A., Cost, A.L., Freikamp, A., Sabass, B., Mehlich, A., Tramier, M., Rief, M., and Grashoff, C. (2017). Multiplexing molecular tension sensors reveals piconewton force gradient across talin-1. Nat Methods 14, 1090–1096.

Rueckert, D., Sonoda, L.I., Hayes, C., Hill, D.L.G., Leach, M.O., and Hawkes, D.J. (1999). Nonrigid registration using free-form deformations: Application to breast MR images. Ieee T Med Imaging 18, 712–721.

Sartori, A., Gatz, R., Beck, F., Rigort, A., Baumeister, W., and Plitzko, J.M. (2007). Correlative microscopy: bridging the gap between fluorescence light microscopy and cryo-electron tomography. J Struct Biol 160, 135–145.

Schaffer, M., Pfeffer, S., Mahamid, J., Kleindiek, S., Laugks, T., Albert, S., Engel, B.D., Rummel, A., Smith, A.J., Baumeister, W., et al. (2019). A cryo-FIB lift-out technique enables molecular-resolution cryo-ET within native Caenorhabditis elegans tissue. Nat Methods 16, 757–762.

Scheres, S.H. (2012). RELION: implementation of a Bayesian approach to cryo-EM structure determination. J Struct Biol 180, 519–530.

Schur, F.K., Obr, M., Hagen, W.J., Wan, W., Jakobi, A.J., Kirkpatrick, J.M., Sachse, C., Krausslich, H.G., and Briggs, J.A. (2016). An atomic model of HIV-1 capsid-SP1 reveals structures regulating assembly and maturation. Science 353, 506–508.

Shemesh, T., Geiger, B., Bershadsky, A.D., and Kozlov, M.M. (2005). Focal adhesions as mechanosensors: a physical mechanism. Proc Natl Acad Sci U S A 102, 12383–12388.

Song, K., Shang, Z., Fu, X., Lou, X., Grigorieff, N., and Nicastro, D. (2019). In situ structure determination at nanometer resolution using TYGRESS. Nat Methods.

Weber, M.S., Wojtynek, M., and Medalia, O. (2019). Cellular and Structural Studies of Eukaryotic Cells by Cryo-Electron Tomography. Cells 8.

Xu, K., Babcock, H.P., and Zhuang, X. (2012). Dual-objective STORM reveals three-dimensional filament organization in the actin cytoskeleton. Nat Methods 9, 185–188.

Zaidel-Bar, R., Itzkovitz, S., Ma’ayan, A., Iyengar, R., and Geiger, B. (2007). Functional atlas of the integrin adhesome. Nat Cell Biol 9, 858–867.

