## Supplemental information and figures for "Revealing the polarity of actin filaments by cryo-electron tomography"

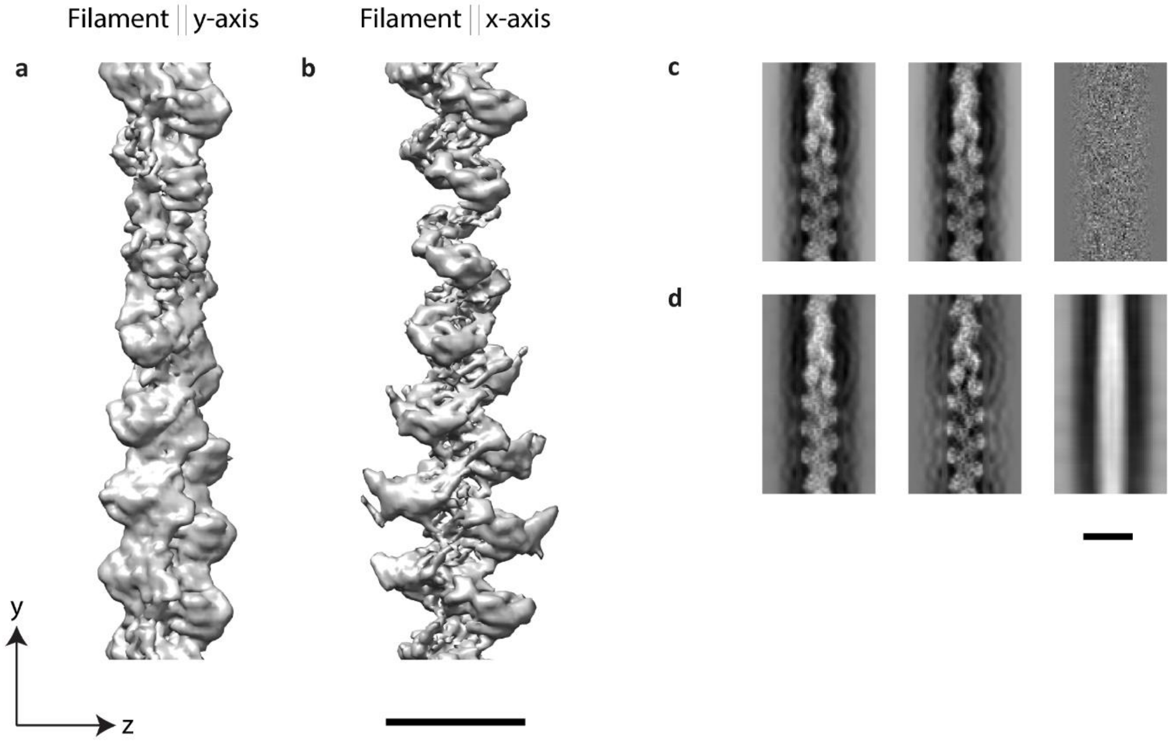


**Supplemental Figure 1 | Missing wedge and projection of subtomograms.** (**a**) Structure of an actin filament (EMD-6179 (Galkin et al., 2015)), that was oriented parallel to the tilt-axis (y-axis), and then distorted by the missing wedge. (**b**) However, if the filament was oriented parallel to the x-axis (orthogonal to the tilt-axis), the anisotropic distortion caused by the missing wedge in z-direction is substantially more pronounced. (**c**) Left to right: projection of filament (**a**) in z-direction, projection of filament (**b**) in z-direction, and difference image between the two projections. The difference image is featureless, which indicates that the missing wedge induced anisotropy vanishes in the projection images. (**d**) However, if a mask in z-direction is applied before projection (in this case the height of the mask was 11 nm), the difference image is not featureless anymore. The influence of this mask on the precision of APT is part of the validation of the method. Scale bars 10 nm.

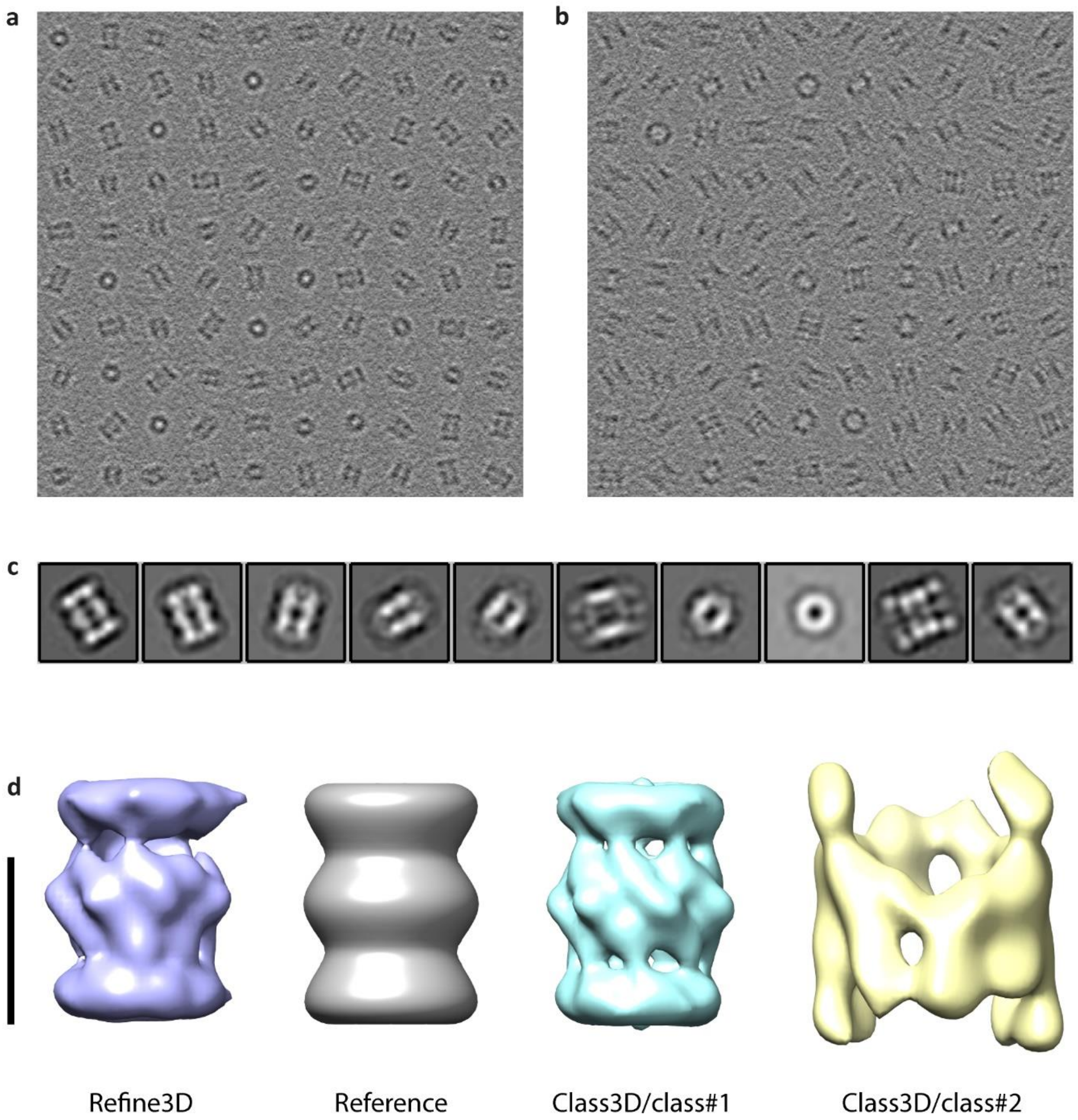


**Supplemental Figure 2 | Classification and structural analysis of projected subtomograms by single particle methods.** We tested our approach for subtomogram averaging with a dataset of modelled subtomograms (Forster et al., 2008), including missing wedge (tilt-range from -60° to +60°), contrast transfer function (defocus of -6 µm, acceleration voltage 300 kV, spherical aberration 2.0 mm, pixelsize 0.42 nm), and modulation transfer function. The dataset contains two particle species: 2048 20S-proteasomes (PDB-1PMA (Lowe et al., 1995)) and 256 thermosomes (PDB-1A6D (Ditzel et al., 1998)). The orientation of the particles was uniformly distributed over the 3D rotation space, and both subtomogram species were modelled with a SNR = 0.1. The subtomograms were projected without masking in the z-direction. (**a**) A gallery of 100 projected proteasomes is displayed. (**b**) A gallery of 100 projected thermosomes is shown. (**c**) The projected images of both particle species were mixed and a 2D classification in RELION (Scheres, 2012) was performed. Since the number of proteasome particles is eight-fold higher than thermosome particles, most of the classes are dominated by proteasomes. (**d**) For 3D reconstruction we initially executed a 3D refinement job in RELION. As starting reference, the proteasome structure was filtered to 45 Å resolution and rotationally symmetrized along the z-axis (grey structure). The obtained reconstruction is shown on the left (blue structure). It resembles a proteasome to a certain degree, but the structure is obviously distorted by the thermosome fraction. Then we performed a 3D classification job in RELION, using the same starting reference as previously, and assuming two classes. Both particle species were properly separated in 3D, with no classification error as compared to the ground truth – all proteasomes were assigned to class#1, cyan structure, and all thermosomes to class#2, yellow structure. The proteasome class average shows the expected D7 symmetry. Scale bar 10 nm.

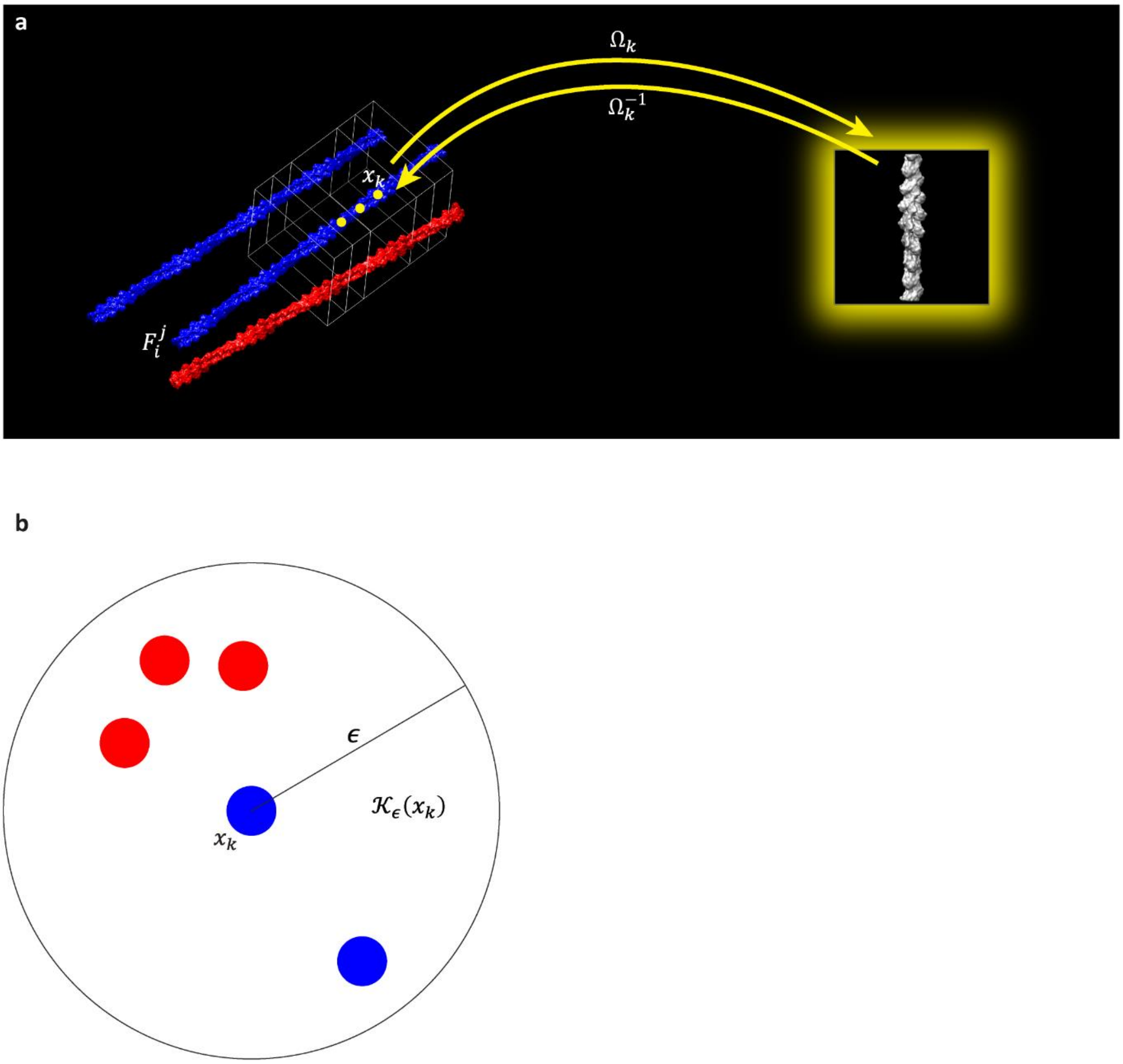


**Supplemental Figure 3 | Basic transformations and topology analysis.** (**a**) The actin filaments $F_i^j$ are subdivided into equidistant segments $x_k$ (yellow dots), each of which is associated with a subtomogram (white boxes). The forward transformation $\Omega_k$ describes how $F_i^j$ has to be transformed at $x_k$ in order to align with the filament average (yellow glowing inset). The inverse transformation $\Omega_k^{-1}$ describes how $F_i^j$ is oriented at $x_k$ with respect to the filament average. Filaments and average were visualized using EMD-6179 (Galkin et al., 2015). (**b**) For topology analysis a sphere $\mathcal{K}_\epsilon(x_k)$ with radius $\epsilon$ is constructed around each $x_k$, and the local polarity distribution within $\mathcal{K}_\epsilon(x_k)$ is analysed. Here $x_k$ is located in a neighbourhood with 75% mixed polarity and 25% uniform polarity, respectively.

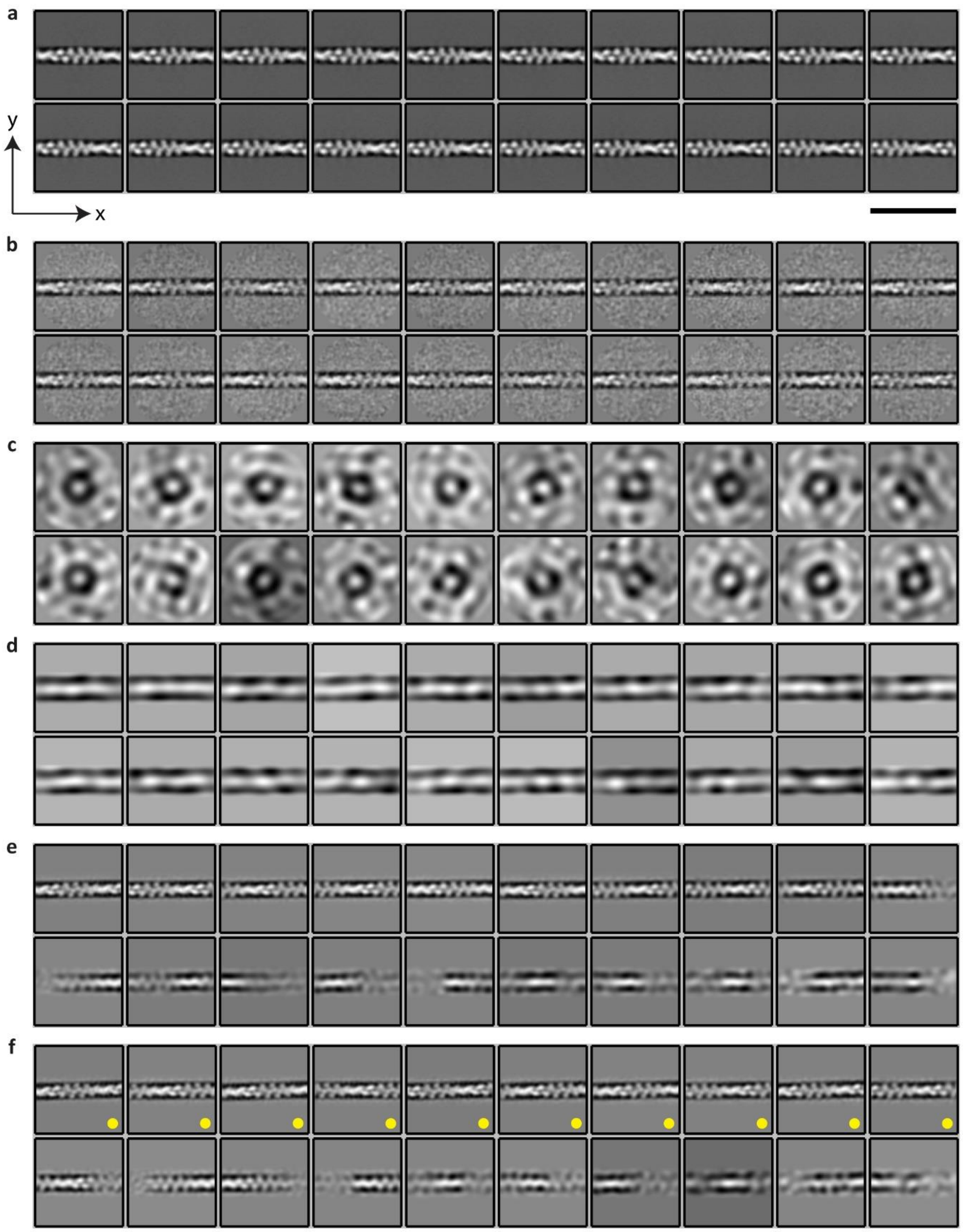


**Supplemental Figure 4 | Prealignment and 2D classification of segments from modelled actin bundles with SNR = 0.0001.** (**a**) The aim of the prealignment step (*Step IV*) is to orient the (central) filament density in the projected subtomograms parallel to the x-axis. Therefore, we used a template library, which was created from an actin filament structure by successively rotating and projecting the filament. The image shows the first twenty entries in the template library. Scale bar 50 nm. (**b**) The first twenty class averages after ten iterations of prealignment are shown. The particle rotations and translations were used as priors for the next step. (**c**) The 2D classification module (*Step V*) of APT aims to produce high quality class averages with RELION (Scheres, 2012). Therefore, we employ the prealignment priors, which allow to apply a mask parallel to the filament during 2D classification. This mask diminishes the influence of neighboring filaments. In contrast to the prealignment step, it is vital that this 2D classification is unsupervised. This ensures that as much as possible structural heterogeneity can cluster in a data-driven way into distinct class averages. The first twenty class averages of the zeroth iteration are shown. It illustrates the initialization of the unsupervised 2D classification. (**d**) In the first iteration the applied mask appears, and the prealignment priors force the central filament density parallel to the x-axis. (**e**) After ten iterations of unsupervised 2D classification the class averages capture most of the structural heterogeneity present in the dataset. (**f**) After 100 iterations we selected the segments, which were assigned to the class averages marked with yellow dots, for subsequent 3D reconstruction (*Step VI*).

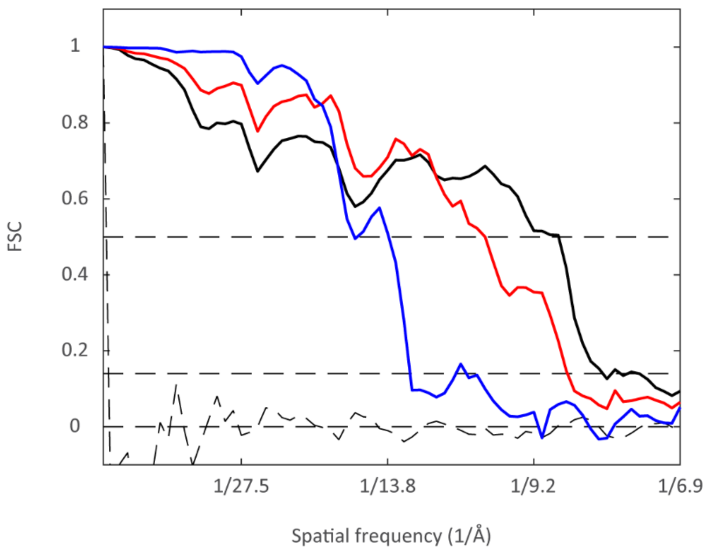


**Supplemental Figure 5 | Resolution measurement of actin filament structures from modelled actin bundles.** Based on the three modelled datasets (SNR=0.01, 0.001, 0.0001), we reconstructed three actin filament structures (~ 10'000 segments per average). Resolution was estimated by Fourier shell correlation (FSC), calculated between the respective averages and EMD-6179 (Galkin et al., 2015), using the 0.5 threshold criterion (Rosenthal and Henderson, 2003). Prior FSC computation, the structures were aligned with each other. As expected, the resolutions of the averages, that are 8.7 Å (black FSC curve, SNR = 0.01), 10.3 Å (red FSC curve, SNR = 0.001), and 15.5 Å (blue FSC curve, SNR = 0.0001), drop with decreasing SNR. The dashed curve is the FSC between two noise volumes, but using an identical masking as for the resolution estimation of the averages. It shows that the mask has no inflating effect on the resolution.

a

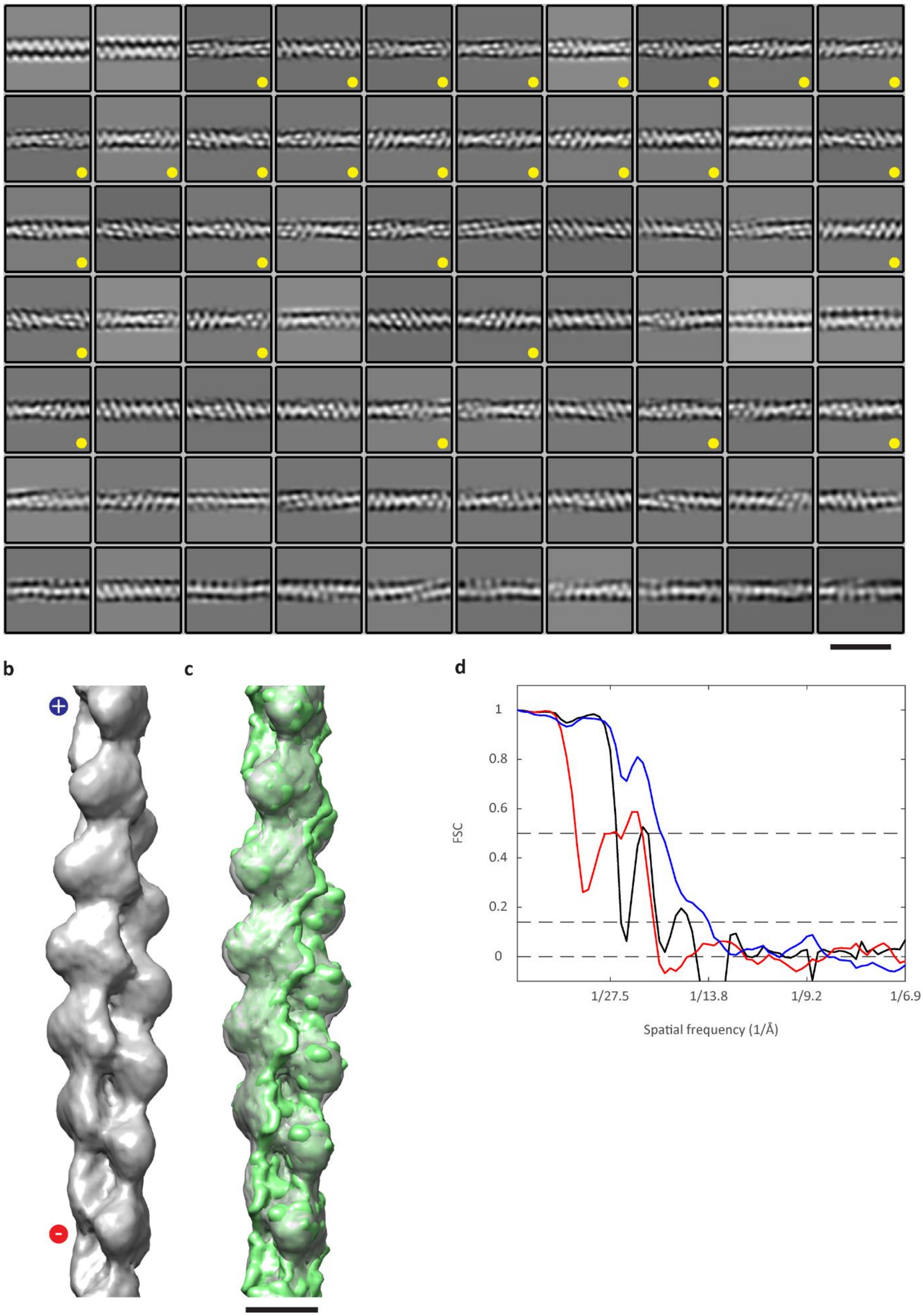

**Supplemental Figure 6 | In-situ actin filament average from manually segmented actin bundles.** (**a**) The image shows the final class averages of the 2D classification (*Step V*). Those segments, which were combined to the class averages marked with yellow dots, were selected for 3D reconstruction (20’585 segments out of 43’400). Scale bar 36 nm. (**b**) In-situ actin filament structure, reconstructed from the selected segments (*Step VI*). The average shows clear polar features, and the position of the plus-end can be detected unambiguously (plus and minus symbols). (**c**) Here we docked EMD-6179 (Galkin et al., 2015) (green isosurface) into our structure (grey isosurface), reaching a correlation value of 0.87. However, if we reverse the filament direction the correlation value drops to 0.73. This shows that the average exhibits polar features. Scale bar 5 nm. (**d**) Resolution was estimated by FSC, calculated between the obtained in-situ average and EMD-6179, serving as an external reference. Prior FSC computation the structures were aligned with each other. The corresponding blue FSC curve crosses the 0.5 threshold criterion (Rosenthal and Henderson, 2003) at 18.4 Å. The black FSC curve was calculated between the 3D reconstruction template and EMD-6179. In comparison with the blue FSC curve it proofs that during 3D reconstruction higher resolution features were successfully extracted from the data (no template bias). The red FSC curve was calculated between the obtained in-situ average and EMD-6179, however, the docking prior FSC computation was conducted with a reversed filament direction compared to the external reference.

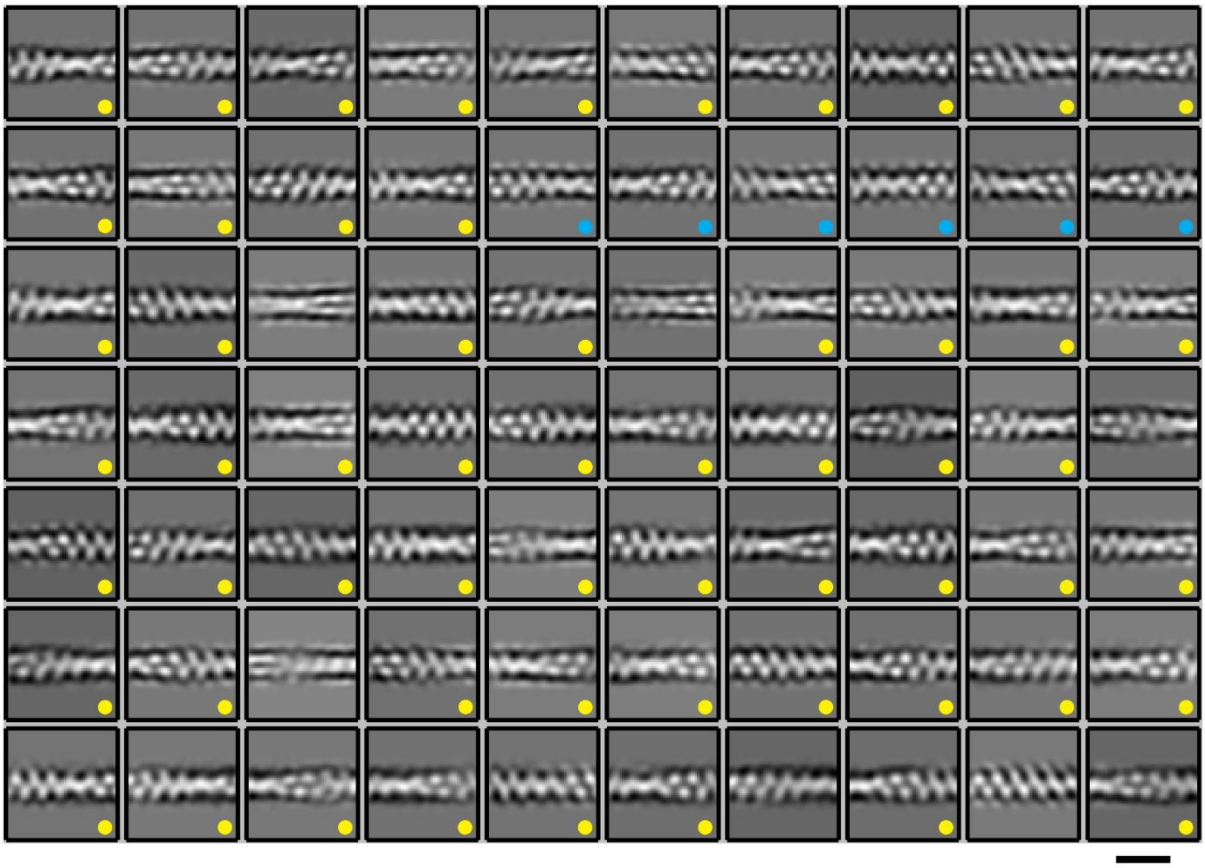

**Supplemental Figure 7 | 2D classification of segments from automatically segmented actin bundles.** The image shows the final class averages (*Step V*). Those segments, which were combined to the class averages marked with yellow dots, were selected for 3D reconstruction (72’973 segments out of 247’940). The class averages marked with blue dots are also shown in Fig. 2d. Scale bar 18 nm.

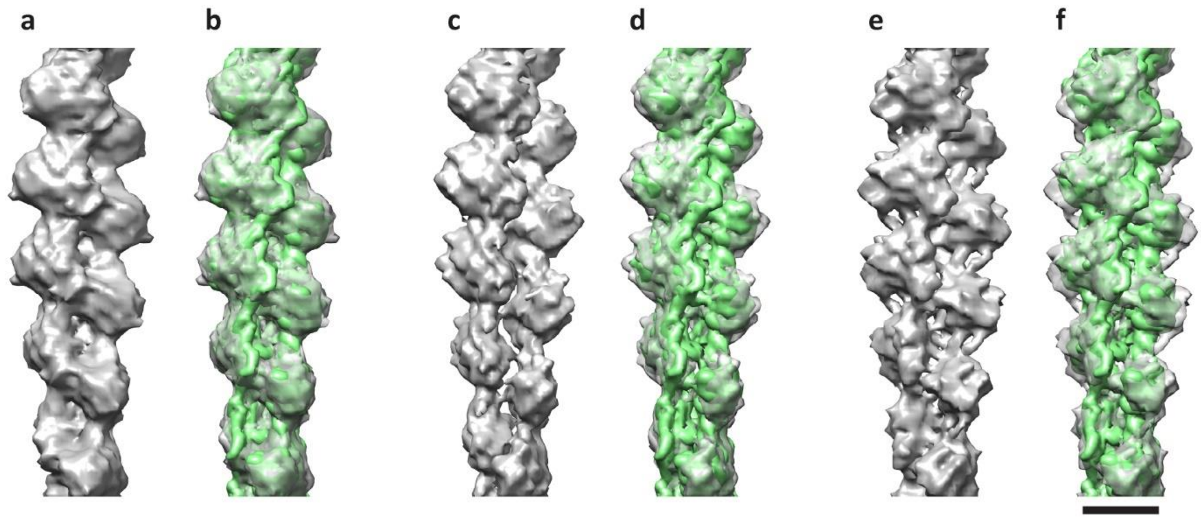


**Supplemental Figure 8 | Structural classes of in-situ actin filaments.** The gallery shows the three class averages (**a**), (**c**), and (**e**) as grey isosurfaces. In (**b**), (**d**), and (**f**) we docked EMD-6179 (Galkin et al., 2015) (green isosurfaces) into the respective structures. Class average (**a**) and docking (**b**) are also displayed in Fig. 2e and Fig. 2f, respectively. Scale bar 5 nm.

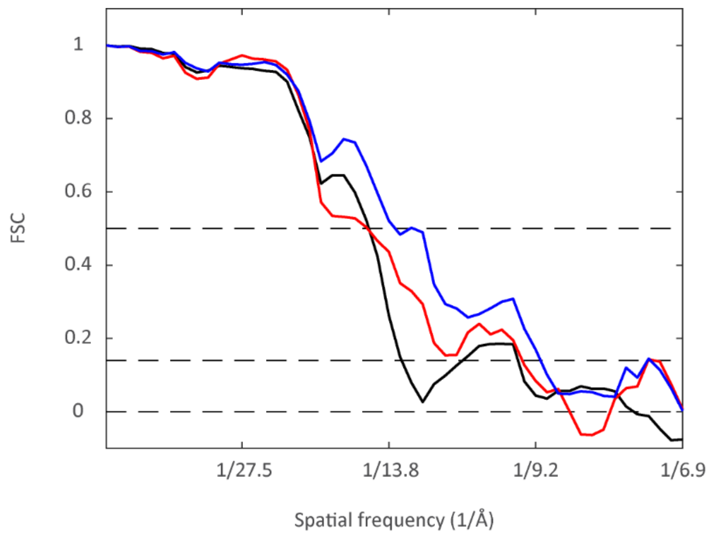


**Supplemental Figure 9 | Resolution measurement of the three class averages obtained from automatically segmented actin bundles.** Resolution was estimated by FSC, calculated between the obtained in-situ class averages and EMD-6179 (Galkin et al., 2015), serving as an external reference. Prior FSC computation the structures were aligned to each other. The corresponding red FSC curve (class#3, Supplementary Fig. 8e), black FSC curve (class#2, Supplementary Fig. 8c), and blue FSC curve (class#1, Supplementary Fig. 8a, Fig. 2e) cross the 0.5 threshold criterion (Rosenthal and Henderson, 2003) at 14.9 Å, 14.8 Å, and 13.5 Å resolution, respectively.

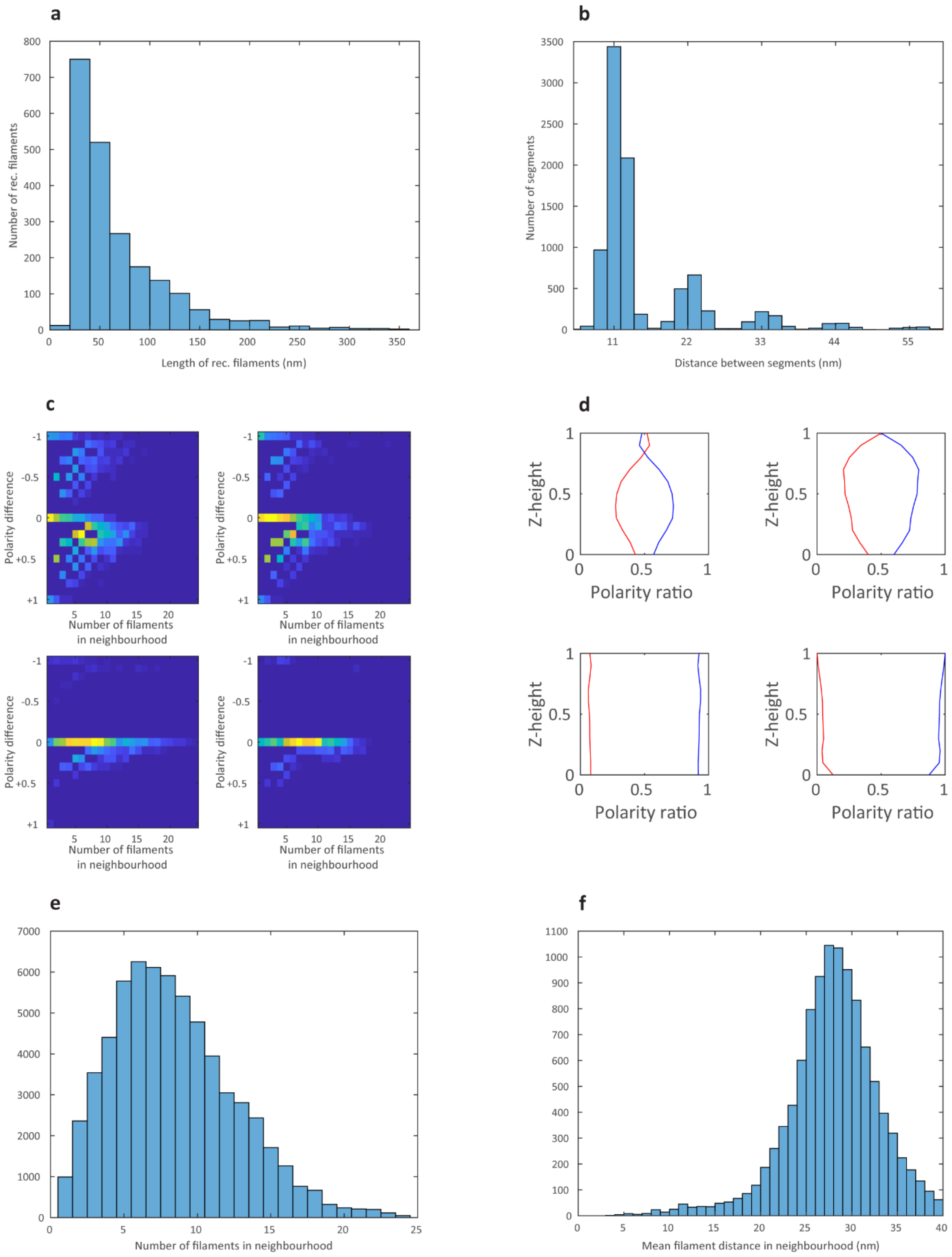


**Supplemental Figure 10 | Analysis of actin bundles at FAs.** (**a**) Length distribution of the reconstructed actin filaments from the manual segmented dataset. (**b**) Initially all segments were extracted with an equidistant spacing of 11 nm. However, during 2D classification (*Step V*) segments were rejected, thus the resulting distance distribution between the segments shows peaks located at multiples of 11 nm. (**c**) Using APT's topology module (*Step IX*) the polarity distributions of the actin bundles were analyzed. Therefore, the depicted matrices were computed based on the bundles shown in Fig. 3e-h (corresponding matrices in the following order: top/left, top/right, bottom/left, and bottom/right). Each matrix element is the number of how often a specific combination of number of filaments within $\mathcal{K}_{\epsilon}(x_k)$ (x-axis) versus the polarity difference within $\mathcal{K}_{\epsilon}(x_k)$ (y-axis) appears in a bundle ($\epsilon$=40 nm). High numbers are visualized as yellow/orange matrix elements, the dark blue background indicates zero. All four bundles show a substantial fraction of uniform polarity (polarity difference = 0). However, only in bundles located at proximal regions of FAs (top/left, top/right) a significant amount of mixed polarity neighborhoods can be detected (polarity difference ≠ 0). (**d**) In order to evaluate where mixed polarity regions are localized in the FA actin bundles, we fitted a plane to the bottom of the bundles (the side which is close to the support), and then stepwise moved this plane upwards through the bundle, and recorded for each step the polarity ratio found within the plane. We executed this calculation for the bundles shown in Fig. 3e-h. The corresponding plots are shown in the following order: top/left, top/right, bottom/left, and bottom/right. The blue trajectory shows the fraction of filaments that are pointing with their plus-ends towards the cell tip at the respective z-height of the plane, which is plotted normalized between 0 (bottom side of the bundle) and 1 (topside of the bundle). The red trajectory shows the fraction of filaments pointing in the opposite direction at respective z-heights. The two distal bundles with predominantly uniform polarity (bottom/left, bottom/right) show a similar polarity ratio at all z-heights. However, in the two proximal bundles (top/left, top/right) a pronounced mixed polarity can be found mainly at bottom and top side of the bundles. (**e**) During topology analysis (*Step IX*) the number of filaments within $\mathcal{K}_{\epsilon}(x_k)$ was evaluated. The histogram shows that the most frequent configuration are six neighboring filaments in the FA actin bundles. (**f**) Additionally, the mean distance of the filaments within $\mathcal{K}_{\epsilon}(x_k)$ was calculated.

### References


Ditzel, L., Lowe, J., Stock, D., Stetter, K.O., Huber, H., Huber, R., and Steinbacher, S. (1998). Crystal structure of the thermosome, the archaeal chaperonin and homolog of CCT. Cell *93*, 125-138.

Forster, F., Pruggnaller, S., Seybert, A., and Frangakis, A.S. (2008). Classification of cryo-electron sub-tomograms using constrained correlation. J Struct Biol *161*, 276-286.

Galkin, V.E., Orlova, A., Vos, M.R., Schroder, G.F., and Egelman, E.H. (2015). Near-atomic resolution for one state of F-actin. Structure *23*, 173-182.

Lowe, J., Stock, D., Jap, R., Zwickl, P., Baumeister, W., and Huber, R. (1995). Crystal-Structure of the 20s Proteasome from the Archaeon T-Acidophilum at 3.4-Angstrom Resolution. Science *268*, 533-539.

Rosenthal, P.B., and Henderson, R. (2003). Optimal determination of particle orientation, absolute hand, and contrast loss in single-particle electron cryomicroscopy. J Mol Biol *333*, 721-745.

Scheres, S.H. (2012). RELION: implementation of a Bayesian approach to cryo-EM structure determination. J Struct Biol *180*, 519-530.
